# Tibetan near-complete pangenome reveals complex variants underlying high-altitude adaptation

**DOI:** 10.64898/2025.12.16.694547

**Authors:** Yaoxi He, Kai Liu, Leyan Mao, Feifei Zhou, Jiaxu Song, Xuebin Qi, Dongya Wu, Junmin Han, Lianting Fu, Jiangguo Li, Yu Zhang, Chaoying Cui, Ouzhuluobu, Guojie Zhang, Yafei Mao, Bing Su

**Affiliations:** State Key Laboratory of Genetic Evolution and Animal Models, Kunming Institute of Zoology, Chinese Academy of Sciences, Kunming, China; Yunnan Key Laboratory of Integrative Anthropology, Kunming, China; University of Chinese Academy of Sciences, Beijing, China; Bio-X Institutes, Key Laboratory for the Genetics of Developmental and Neuropsychiatric Disorders, Ministry of Education, Shanghai Jiao Tong University, Shanghai, China; State Key Laboratory of Primate Biomedical Research Institute of Primate Translational Medicine, Kunming University of Science and Technology, Kunming, China; High Altitude Health and Translational Medicine Institute, Tibetan Fukang Hospital, Lhasa, China; Women’s Hospital, Zhejiang University School of Medicine & Center for Evolutionary & Organismal Biology, Zhejiang University, Hangzhou, China; International Institutes of Medicine & the Fourth Affiliated Hospital, Zhejiang University School of Medicine, Yiwu, China; Institute of Bioinformatics & James D. Watson Institute of Genome Sciences, Zhejiang University, Hangzhou, China; Xizang Fukang Hospital, Lhasa, China; Liangzhu Laboratory, Zhejiang University Medical Center, Hangzhou, China; Center for Excellence in Animal Evolution and Genetics, Chinese Academy of Sciences, Kunming, China

## Abstract

Genomic diversity in indigenous populations offers critical insights into human evolution and population-specific adaptation. Here, we generated 70 near-complete, fully phased genome assemblies from 35 high-altitude Tibetan trios and identified 4.55 million small variants and 63,031 structural variants (SVs) absent from the current global pangenome reference. Comparative analyses with a 595 globally diverse long-read genome dataset revealed 207 previously uncharacterized Tibetan-enriched SVs spanning 275 genes implicated in high-altitude physiological adaptation. Genotyping these SVs in 1,164 Tibetan and Han individuals with 88 physiological traits reveals five variants significantly associated with cardiopulmonary and hematological phenotypes central to hypoxia adaptation. We further map 2.45 Gb of archaic hominin sequences in Tibetans and uncover 18 previously unrecognized introgressed regions enriched in Tibetans with genes related to hematopoiesis and lung function. These regions harbor five Tibetan-enriched SVs, indicating that archaic introgression contributed SVs relevant to human adaptation. Together, these findings reveal substantial uncharacterized genomic diversity in Tibetans and highlight a previously underappreciated role of SVs and archaic SVs in shaping human evolution and high-altitude adaptation.

## Introduction

Deciphering every nucleotide in a diploid human genome is fundamental to elucidating the genetic underpinnings of phenotypes, diseases, genome evolution, and local adaptation. Although traditional short-read sequencing has enabled broad surveys of human genetic diversity, its intrinsic limitations—particularly in resolving large, repetitive, and structurally complex regions—have left substantial portions of the genome inaccessible to comprehensive analysis (Logsdon et al., 2025). Recent advances in telomere-to-telomere (T2T) genome assembly and pangenome reference construction are transforming this landscape. These technologies now enable near-complete reconstruction of human genomes and capture population-level genomic diversity with unprecedented resolution. Collectively, they recover approximately 237 Mbp (∼8%) of previously unresolved sequence, encompassing highly repetitive and structurally complex regions that harbor extensive and biologically meaningful genetic variation (Nurk et al., 2022). Yet despite substantial progress in T2T and pangenome research, how complex genomic variants contribute to human evolutionary adaptation remains largely unknown.

High-altitude adaptation in Tibetans offers a powerful natural model for examining how genomic variation shapes human adaptation. Extensive phenotypic studies have shown that Tibetan adaptation involves multiple physiological systems, including hematological traits (e.g., a blunted erythropoietic response), cardiopulmonary function, vascular regulation, metabolic processes, and reproduction (Beall, 2007; He et al., 2023a; He et al., 2023b; Wu and Kayser, 2006). Genomic studies, primarily focused on single-nucleotide variants (SNVs), have identified several key loci, including *EPAS1*, *EGLN1*, and *PPARA*, associated with hemoglobin concentration, red blood cell regulation, and hypoxia signaling (Beall et al., 2010; Peng et al., 2011; Simonson et al., 2010; Xiang et al., 2013; Yi et al., 2010). Earlier efforts to characterize structural variation in Tibetans using long-read sequencing (He et al., 2020; Quan et al., 2021; Shi et al., 2023) have provided valuable early insights but were constrained by low-coverage data and the inability to resolve structurally complex regions, including peri/centromeres, rDNA arrays, and long tandem repeats. Moreover, the absence of deep phenotypic measurements in most datasets has limited the interpretation of whether identified complex variants influence altitude-related traits.

Additionally, Tibetan genomes harbor a Denisovan-derived haplotype at the *EPAS1* locus (Huerta-Sánchez et al., 2014; Zhang et al., 2021), which confers advantageous physiological responses to chronic hypoxia. This remarkable example of archaic introgression highlights the complex evolutionary processes that have contributed to Tibetan adaptation. However, the full extent of archaic-derived genomic content in Tibetans, especially within previously unresolved genomic regions, is still unknown.

Here, we address these limitations by generating the most comprehensive genomic resource to date for an indigenous high-altitude human population. We sequenced 35 Tibetan trios and produced the first T2T level pangenome graph specifically tailored to a high-altitude population. This resource provides an unprecedented view of the full spectrum of Tibetan genomic diversity and high-altitude adaptation, including sequences absent from current global pangenome references, complex structural variants, and archaic introgressed fragments with base-level precision. Collectively, this study provides a representative framework for exploring evolutionary history and local adaptation using T2T and pangenome technologies, and it opens new avenues for dissecting the genomic architecture underlying complex traits in diverse human populations.

## Results

### Assembling 35 fully phased Tibetan genomes from trio samples

We collected 105 indigenous Tibetans (35 trios) from five geological locations of Xizang Autonomous Region of China (**Fig. 1a; Supplementary Fig. 1a; Supplementary Table 1**). Using these samples, by integrating multiple deep-sequencing data for each individual, we generated 70 fully phased assemblies of Tibetans, including eight near-complete and 27 high-quality Tibetan diploid genomes (**Methods**). On average, each near-T2T assembly was supported by 50× PacBio HiFi data, 95× ultra-long Oxford Nanopore Technology (ONT) data, 38× short-read sequencing data, and 66× Hi-C data (**Fig. 1b; Supplementary Fig. 1b; Supplementary Table 2**).

**Figure 1.**
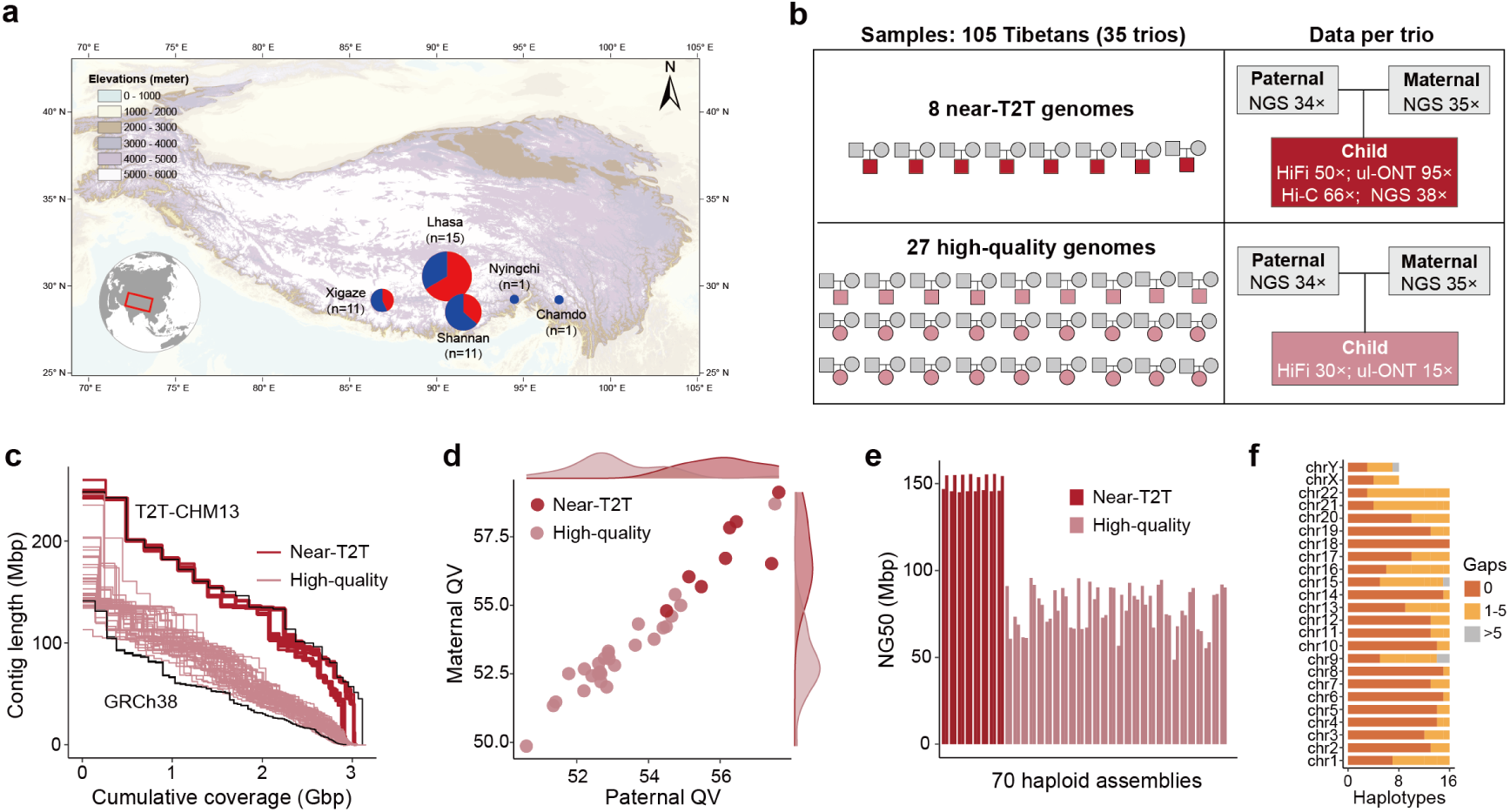
Sampling, data generation, and assembly statistics. **a**, Five sampling sites on the Tibetan Plateau are indicated. The size and color of each pie chart represent sample size and sex composition (red, male; blue, female), respectively. Elevation (meters) is indicated by the color bar.,. The elevations of the sampling sites are 3,253∼5,020 meters above the sea level. The map template was generated using ArcGIS. **b,** Overview of sample collection and data generation. A total of 105 Tibetan individuals (35 complete trios) were collected to assemble eight near-T2T genomes and 27 high-quality genomes using multiple sequencing technologies. T2T, telomere-to-telomere; NGS, next-generation sequencing; HiFi, PacBio high-fidelity (HiFi) long-read sequencing; ul-ONT, Oxford Nanopore Technologies ultra-long-read sequencing; Hi-C, high-throughput chromosome conformation capture sequencing. **c,** Line plot showing the assembly contiguity (contig sizes) of the 70 haploid assemblies. Contigs of two reference genomes (T2T-CHM13 and GRCh38) are shown for comparison. **d,** Assembly QVs indicating base-level accuracy for maternal and paternal assemblies of each trio. **e,** Contig NG50 values of the 70 haploid assemblies. For each sample, the left bar represents the paternal haplotype and the right bar the maternal haplotype. The eight near-T2T and 27 high-quality samples are shown in dark and light red, respectively. NG50 is defined as the contig length at which half of the predicted genome size is contained in contigs at least that length. **f,** Chromosome counts for the 16 near-T2T haploid assemblies under three gap-count thresholds: gapless, 1–5 gaps, and >5 gaps.

We assessed the genome completeness, contiguity and accuracy for all 70 haploid assemblies (**Figure 1c-e**; **Supplementary Fig. 1c,1d; Supplementary Table 3**). The 16 near-T2T haploid assemblies had an average genome size of 3.06 giga base pairs (Gbp) (3.04–3.09 Gbp), an average quality value (QV) of 56.5 (55.0–57.9), and an average contig N50 of 152.5 Mbp (149.2–151.2 Mbp). Among them, 100 chromosomes (56%) were fully assembled without gaps, and 230 chromosomes (95%) contain ≤5 gaps (**Fig. 1c-f**). On average, 99.2% (98.6–99.6%) of sequences in each assembly were correctly assembled according to Flagger evaluation (Liao et al., 2023) (**Supplementary Fig. 1e**)

For the remaining 54 high-quality assemblies, the average genome size was 3.01 Gbp (3.00–3.02 Gbp), with an average QV of 53.5 (50.0–55.9) and an average contig N50 of 80.5 Mbp (60.2–90.2 Mbp) (**Fig. 1c-e**). Flagger analysis showed that an average of 93.0% (90.7–94.9%) of sequences were correctly assembled (**Supplementary Fig. 1d)**.

### Structural landscape of telomeres and centromeres in Tibetans

To investigate previously unresolved regions in Tibetan genome, we analyzed telomeres, centromeres and complete chromosome Y using 16 near-T2T Tibetan haploid assemblies (**Methods**). Across these assemblies, we successfully reconstructed telomeres for 364 chromosomes, representing 98.91% of all chromosomes analyzed (**Supplementary Fig. 2a; Supplementary Table 4**). Among them, 278 chromosomes (75.54%) contained telomeres at both ends, while 86 chromosomes (23.37%) contained a telomere at one end (**Supplementary Fig. 2b**) Notably, chromosomes 13, 14, 15, and 21 were disproportionately represented among the one-ended assemblies, consistent with previous reports that these acrocentric chromosomes are enriched for rDNA arrays and macrosatellite repeats, making them particularly challenging to assemble (de Lima et al., 2025; Nurk et al., 2022) (**Supplementary Fig. 2c**). Four chromosomes lacked assembled telomeres at both ends, including chromosomes 5, 14, 15, and 19 in four assemblies (**Supplementary Fig. 2c**). The total telomere length per individual averaged 261.0 kbp (231.1–292.7 kbp) **(Supplementary Table 4)**, with no significant difference in telomere length among chromosomes (*p* = 0.22, Dunn’s test) **(Supplementary Fig. 2d)**

Sequence identity analyses revealed substantial structural diversity in Tibetan telomeres (**Supplementary Fig. 3a**). Using a statistical classification model (**Methods**), we categorized telomeres into three structural types: Continuous (C), Fragmented (F), and Blocky (B) (**Fig. 2a; Supplementary Table 5**). C-type telomeres showed minimal or no mutations within the canonical TTAGGG motif; F-type telomeres harbored random mutations in TTAGGG motif, while B-type telomeres contained recurrent, motif-specific substitutions. Among them, C-type telomeres were the most prevalent (78.40%), followed by B-type (13.93%) and F-type (7.67%) (**Fig. 2b**). For B-type telomeres, substitutions primarily involved T>C or T>G mutations at the first two nucleotides of the TTAGGG repeat, forming four subtypes. Among these, the T(T>G)AGGG pattern was the most common, accounting for 49.44% of all B-type telomeres (**Fig. 2c**). This pattern was also observed in other global populations (**Supplementary Fig. 3b**).

**Figure 2.**
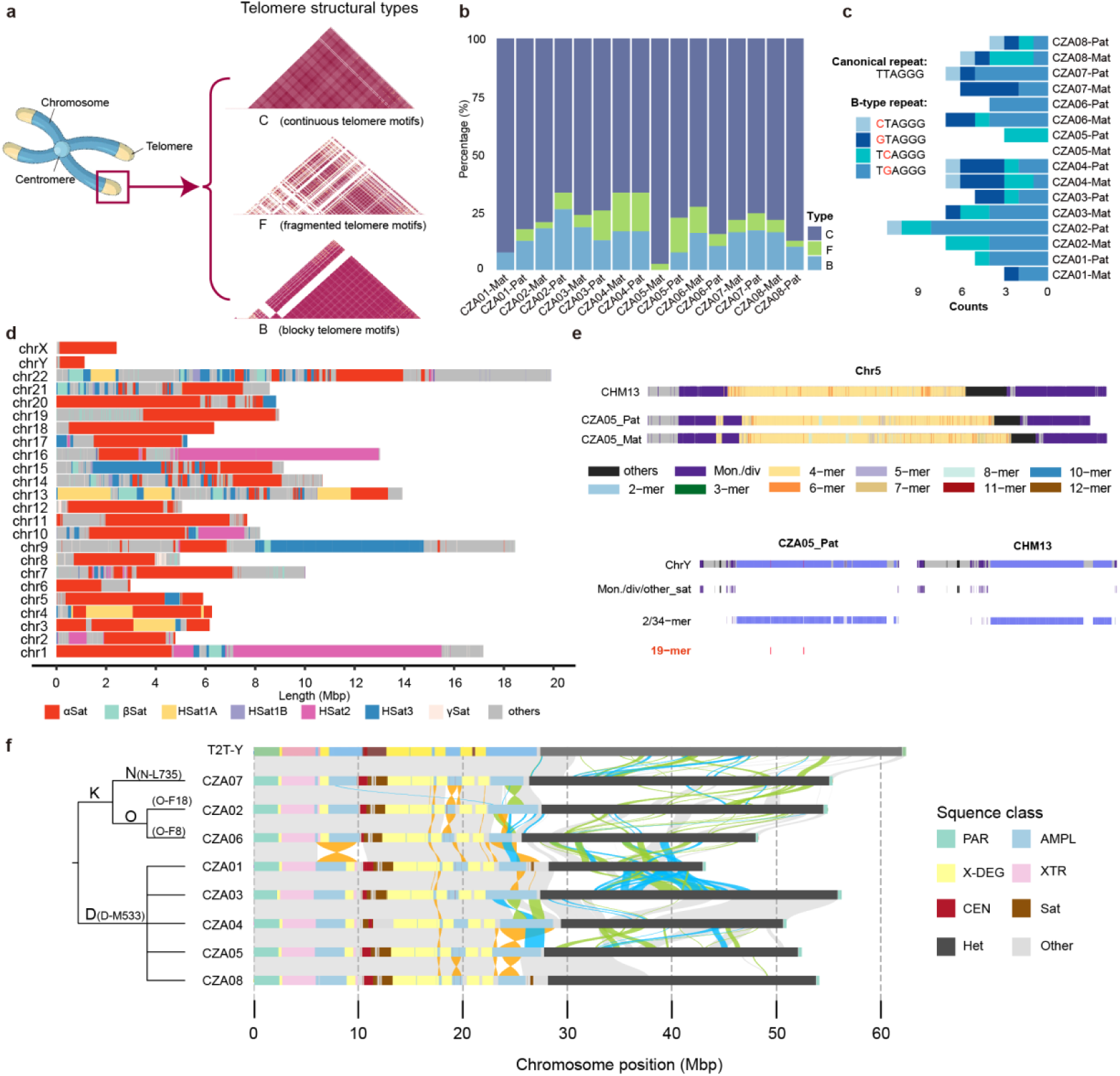
Characterization of Tibetan telomeres, centromeres, and chromosome Y. **a**, Structural diversity of Tibetan telomeres. Telomeres were classified into three categories based on sequence identity: Continuous (C), Fragmented (F), and Blocky (B). **b,** Proportion of three telomere types across all haploid assemblies. **c,** Counts of the four motif variants observed in B-type telomeres. The canonical telomeric repeat and the four mutated motif variants are shown. **d,** Color-coded map of peri/centromeric satellite DNA arrays across all Tibetan assemblies. The colored blocks represent different satellite arrays, including α-satellite (αSat), β-satellite (βSat), H-satellite1A (HSat1A), H-satellite1B (HSat1B), H-satellite2A (HSat2A), H-satellite3 (HSat3), γ-satellite (γSat). **e,** Comparison of α-satellite higher-order repeat (HOR) arrays on chromosome 5 and chromosome Y between Tibetan assemblies and CHM13. HORs are colored according to the number of α-satellite monomers in each repeat unit. **f,** Phylogenetic relationships (left) and haplogroup assignments of the analyzed Tibetan Y chromosomes. Alignment between CZA-Y and T2T-Y is shown on the right; regions with >95% sequence identity are linked and colored by alignment orientation (gray, syntenic; orange, inversion; green, translocation; blue, duplication). AMPL, ampliconic; CEN, centromere; Sat, satellites; Het, heterochromatic.

We next annotated centromeric regions across all chromosomes, totaling 209.6 Mbp (range: 177.8–223.7 Mbp), corresponding to 7.0% (6.1%–7.4%) of the genome (**Fig. 2d** and **Supplementary Fig. 4**). Centromeres on the same chromosome across different individuals showed relatively little variation in size and structure. These regions consisted mainly of four major satellite families: α-satellite, β-satellite, γ-satellite and H-satellite (**Supplementary Fig. 5a**). Considering α-satellite arrays form the core functional component of the human centromere, and account for an average of 40.4% of each Tibetan centromere (**Supplementary Table 6**), we performed an in-depth analysis of the α-satellite arrays in Tibetans. We measured the variation in the length of the higher order repeat (HOR) arrays in centromere α-satellite on each chromosome (**Supplementary Table 7**). The average α-satellite HOR array length was 3.3 Mbp and varied significantly across chromosomes (0.2–8.9 Mbp) (*p* < 2.9×10^-11^, Dunn test) (**Supplementary Fig. 5b**). To identify Tibetan-specific HOR structure, we compared the centromeric sequences from 16 Tibetan assemblies with those from 152 diverse Asian genomes in the Asian Pangenome phase 1 (APGp1) (Dongya Wu et al.). We identified two Tibetan-specific HOR arrays (**Fig. 2e**), including one on chromosome 5 where a monomeric α-satellite insertion disrupted the canonical D1Z7 HOR array, splitting it into two distinct arrays, and the other one on chromosome Y where the organization of the novel α-satellite HOR array was distinct from that of the APGp1 and T2T-CHM13 arrays (**Fig. 2e**).

### Phylogenetic and structural evolution of Tibetan Y chromosome

Human Y chromosome contains extensive repetitive sequences, which historically hindered complete assembly and limited its genomic characterization. Here, we generated eight complete Tibetan Y chromosomes (**Supplementary Table 8**). Phylogenetic comparisons showed strong concordance with the T2T-Y reference, indicating high assembly accuracy. Tibetan Y chromosomes exhibited substantial structural and size variation (Figure 2f), ranging from 43.1 to 56.0 Mbp. The heterochromatic Yq12 region showed the greatest variability (14.7 to 28.7 Mbp, mean = 24.2 Mbp), whereas euchromatic regions were more conserved, except for pronounced variation in ampliconic subregion 2 (**Figure 2e**).

We investigated the variants in Tibetan Y chromosome using DeepVariant (Yun et al., 2021) and PAV (Ebert et al., 2021) (**Methods**). In the euchromatic region, each assembly contained an average of 338 insertions/deletions (≥50 bp), three large inversions (>1 kbp), 2,530 indels (<50 bp), and 10,045 SNVs relative to the T2T-Y reference (Rhie et al., 2023) (**Supplementary Fig. 6a**). On average, each Tibetan Y chromosome harbored six large inversions, four of which overlapped variants identified by sequential variant calling. Notably, we identified a 4.2 Mbp inversion spanning the X-transposed (XTR), ampliconic (AMPL), and X-degenerate (X-DEG) regions, with a strikingly high frequency in the D-M533 lineage, a Tibetan-specific Y chromosome lineage (Figure 2f). This inversion was absent in other Tibetan lineages, even after expanding our dataset to 17 Tibetan male assemblies (**Supplementary Fig. 6b**). Comparative analyses with APG and HPRC assemblies showed that this inversion is present across multiple global lineages, including D, E, C, R, O, and J (**Supplementary Fig. 6c**), suggesting that this region is likely a hotspot for inversions in human Y chromosomes.

### Pangenome reference graph of Tibetan population

Using Minigraph-Cactus pipeline (**Methods**) (Hickey et al., 2024), we constructed a graph-based pangenome of the Tibetan population, incorporating 70 haploid assemblies anchored to the T2T-CHM13 . The resulting graph spans 3,241,333,356 bp in total length. We identified 122.05 Mbp of non-reference sequences, including 34.42 Mbp of singleton sequences. Notably, 65.03 Mbp non-reference sequences are common (present >5% in all haplotypes), representing previously unsolved core genomic regions in Tibetan population (**Fig. 3a**).

**Figure 3.**
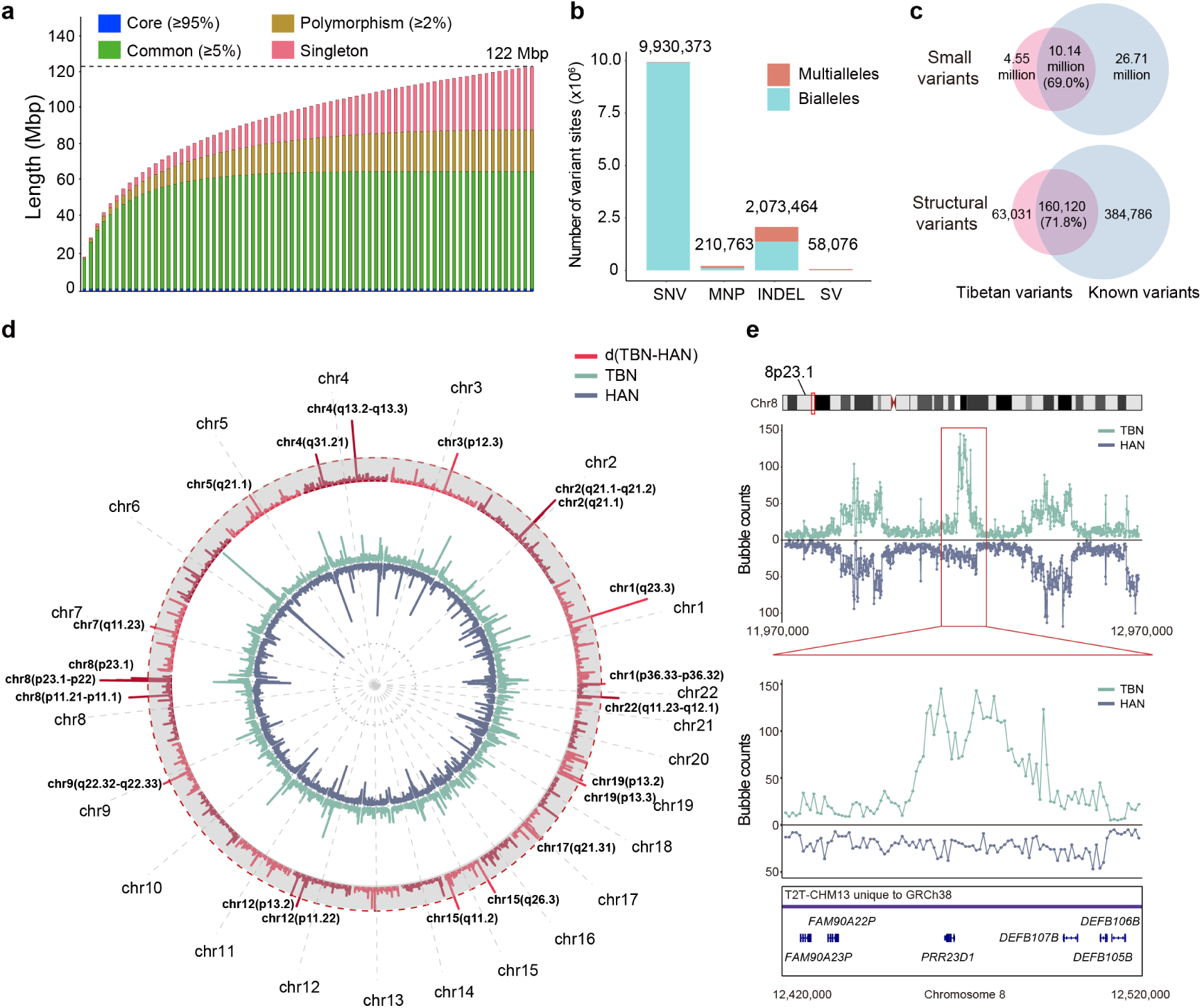
Pangenome graph of Tibetan population. **a,** Cumulative growth curves of the 35-individual Tibetan pangenome graph. The bar height indicates how often each non-reference segment appears across assembled haplotypes. Non-reference sequences were classified into four categories based on their frequency: core (≥95%), common (≥5% and <95%), polymorphic (≥2% and <5%), and singleton (present in only one haplotype). **b,** Variant sites identified in the pangenome graphs, stratified by variant type and allele counts per site. MNP, multi-nucleotide polymorphism. **c,** Number of Tibetan-specific and shared variants between the Tibetan set and the known sets (union of HPRC and HGSVC3) derived from the pangenome graph. **d,** Genome-wide bubble counts of Tibetans (TBN), Han Chinese (HAN), and their difference (d(TBN-HAN)) calculated in 1-Mbp sliding windows. The red dashed line marks the top 0.1% of d(TBN-HAN) values. The 27 genomic regions with significantly elevated bubble counts in Tibetans are labeled. **e,** The most differentiated genomic region in view of bubble counts between Tibetan and Han Chinese populations at 8p23.1 is shown, with a 1 Mbp window displayed in the upper panel and a zoomed-in 100 kbp window in the lower panel. Each dot represents the bubble count within a 10 kbp sliding window for Tibetans (TBN, shown in green) and Han Chinese (HAN, shown in blue).

Across the Tibetan pangenome, we detected 12,272,676 variant sites, comprising 9.9 million SNVs, 210,763 multi-nucleotide polymorphisms (MNPs), 2.1 million indels and 58,076 structural variants (**Supplementary Table 9**). Among these, 93% were biallelic and 7% were multiallelic (**Fig. 3b**). Each sample contained an average of 4.59 million small variants (<50 bp) (range: 4.54–4.68 million), and each haplotype harbored an average of 14,557 SVs (≥50 bp) (**Supplementary Fig. 7a**). We benchmarked variants decoded from the pangenome graph against those identified using conventional reference-based genotyping approaches (DeepVariant, PBSV, Sniffles2, SVIM-asm, and PAV) (**Methods**). Overall, we observed strong concordance between the graph-based and reference-based variant sets across variant classes (**Supplementary Fig. 7b**), with mean recall and precision of 96.96% and 99.20%, respectively, for small variants in non-complex regions (∼85.8% of the autosomal genome). Performance for SVs was slightly lower (94.32% recall and 93.05% precision) (**Supplementary Table 10**). The SV length distribution showed clear peaks at ∼300 bp (Alu insertions) and ∼6 kbp (LINE-1), further supporting the reliability of the pangenome-derived variant calls (**Supplementary Fig. 7c**). Taken together, the variant set derived from the Tibetan graph is of high quality and suitable for downstream analyses.

To assess novelty, we compared our variant set to the union of variants reported by the Human Genome Structural Variation Consortium Phase 3 (HGSVCp3) (Logsdon et al., 2025) and the Human Pangenome Reference Consortium Phase 1 (HPRCp1) (Liao et al., 2023) (**Methods**). We found that 4.55 million small variants (31.0%) and 63,031 SVs (28.2%) were absent from the combined known set (**Fig. 3c**), suggesting that these represent previously unreported variation in the Tibetan population.

We next examined population-stratified genomic regions by comparing bubble counts in the Tibetan pangenome graph with those in a Han Chinese graph based on APGp1 samples (**Methods**). In total, we identified 27 regions with significantly elevated bubble counts in Tibetans (Wilcoxon test, *p* < 2.2 × 10⁻¹⁶; **Supplementary Table 11**). Of these regions, five correspond to sequences unique in T2T-CHM13 compared to GRCh38, and eleven contain protein-coding genes (**Supplementary Fig. 8**). Notably, eight of the twenty-seven regions overlap with Neanderthal- or Denisovan-derived introgressed haplotypes (**Supplementary Fig. 8**), suggesting that archaic introgression may be an important source contributing to the elevated genomic diversity observed in these Tibetan-specific regions.

One notable example was a 1-Mbp region containing a cluster of defensin (DEF) genes at 8p23.1 (chr8: 11,970,000–12,970,000; T2T-CHM13 coordinates), a highly dynamic and structurally complex locus in current human reference genomes. This region exhibited pronounced Tibetan-specific diversity (**Fig. 3e**). The DEF genes (*DEFB105/106/107B*) encode key components of the innate immune system that protect against bacterial and viral infections and can even trigger antitumor immune responses (Taudien et al., 2004).

### Copy number variation of protein coding genes

We annotated copy number variations (CNVs) of protein-coding genes in the Tibetan pangenome. On average, 92.4% of T2T-CHM13 protein-coding genes were covered per assembly, and 333 genes per assembly (excluding sex chromosomes) contained CNVs, including 51 copy-gain and 282 copy-loss genes (**Supplementary Table 12**).

We then compared gene copy numbers between Tibetans and lowland populations using Tibetan assemblies together with reference genomes from APGp1, HPRCy1, and HGSVC3. In total, 44 protein-coding genes exhibited significant copy-number differences between Tibetans and lowlanders, defined as a frequency difference >0.2 for at least one copy state (**Fig. 4a; Supplementary Table 13**). The most divergent gene is *APOBEC3B*, which exhibits a markedly higher frequency of gene loss in Tibetans than in non-Tibetan populations (55% *versus* 20%; **Fig. 4b**). Previous studies have shown that *APOBEC3B* deletion is more common in East Asians (27%) than in Europeans (5%) or Africans (12%) (Kidd et al., 2007), and it plays an important role in innate antiviral immunity by inhibiting retroviral infection (An et al., 2009), and is associated with an increased risk of breast cancer (Han et al., 2016). We also identified a population-stratified CNV at *PHACTR1*, which shows a substantially higher frequency of gene loss in Tibetans compared with non-Tibetans (40% *versus* 15%; **Fig. 4b**). *PHACTR1* is involved in endothelial cell survival and tubule formation, and its genetic variation has been linked to susceptibility to hypertension (Fujimaki et al., 2015), myocardial infarction, coronary artery disease, and cervical artery dissection (Debette et al., 2015).

**Figure 4.**
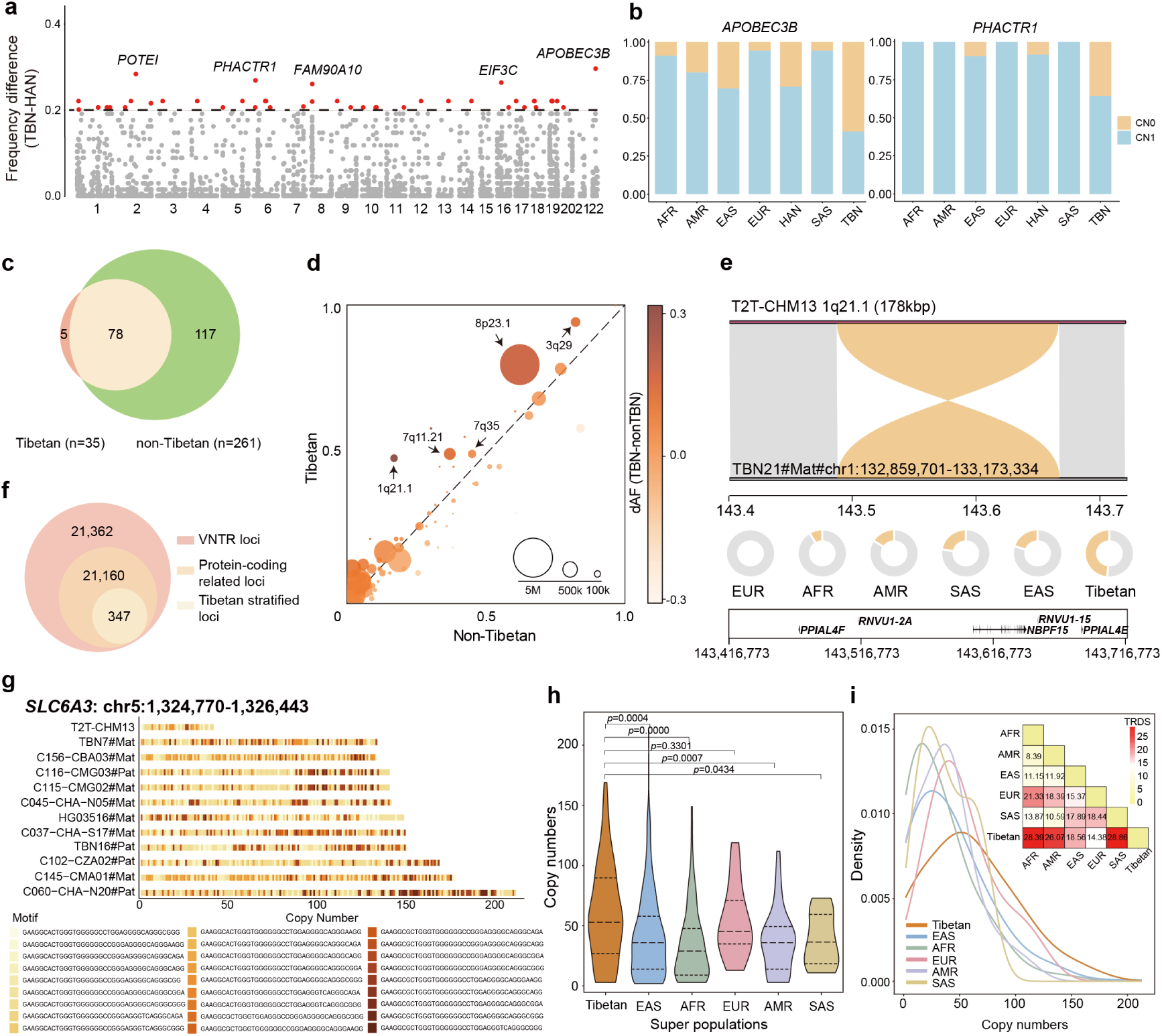
CNV, inversion and VNTR diversity in Tibetan individuals. **a**, Genome-wide frequency differences of copy number variations (CNVs) in protein-coding genes between Tibetans (TBN) and Han Chinese (HAN). Genes with highly divergent copy number frequencies (difference > 0.2) are highlighted in red, and the top five signals are labeled. **b,** Proportions of gene copy numbers across populations. CN0, copy number = 0; CN1, copy number = 1. AFR-African, AMR-American, EAS-East Asian, EUR-European, SAS-South Asian. **c,** Venn diagram showing the overlap between inversions identified in the Tibetan (n = 35) and non-Tibetan genomes (APGp1 callset, n = 261). **d,** Comparison of allele frequencies of large inversions (≥10 kbp) between Tibetan and non-Tibetan populations. Differences of allele frequency (dAF) highlight population stratification, with notable loci at 1q21.1, 8p23.1, 3q29, 7q11.21, and 7q35 showing Tibetan-elevated frequencies. **e,** Synteny plot and population frequency distribution of the 1q21.1 inversion (∼178 kbp) across six global super-populations. EUR, European; AFR, African; AMR, American; SAS, South Asian; EAS, East Asian. **f,** VNTR loci identified in this study include 347 Tibetan-stratified VNTRs within protein-coding regions (total VNTRs: 42,769; protein-coding-related: 21,507; Tibetan-stratified: 347). **g,** A VNTR locus within an intron of the *SLC6A3* gene shows individual-level copy number variation. **h,** Copy number distribution of the VNTR in Tibetans *versus* other super-populations. Statistical significance was assessed by Mann–Whitney U test. **g,** Density plot of the *SLC6A3*-associated VNTR copy number across six super-populations, with Tibetans (orange) showing a significantly higher copy number. Heatmap shows pairwise 2-Wasserstein distances (Tandem Repeat Disparity Score, TRDS) between populations; color scale from 0 (yellow, identical) to 30 (red, maximal divergence).

### Complex structural variants in Tibetan genome

We systematically characterized large genomic inversions (≥10 kbp) in 35 unrelated Tibetan individuals using three tools—PAV (Ebert et al., 2021), PBSV (https://github.com/PacificBiosciences/pbsv), and LSGvar (Zhang et al., 2024) — and identified 83 high-confidence inversions. Cross-referencing with the APGp1 inversion callset (n = 261), showed that 93.98% (78/83) were concordantly detected, with five representing Tibetan-stratified variants, including novel events at 4q13.3 and 15q13.2 (**Fig. 4c; Supplementary Table 14; Supplementary Fig. 9a**).

Population-level comparison revealed 23 inversions with significant Tibetan-specific frequency shifts relative to at least two global super-populations (EAS, AFR, EUR, AMR, and SAS; Fisher’s exact test, *p* < 0.05). Of these, 11 inversions showed elevated allele frequencies in Tibetans, several within regions previously associated with genomic instability (e.g., 8p23.1, 7q11.21, and 3q29; **Figure 4d; Supplementary Figure 9b**) (Wang et al., 2023). Notably, a ∼178 kbp inversion at 1q21.1 (T2T-CHM13, chr1:143,494,155–143,672,549), previously linked to developmental phenotypes (Mefford et al., 2008), was observed in 48.6% of Tibetan genomes compared to ∼20% of East Asian genomes (n = 167), suggesting population-specific selection or genetic drift (**Fig. 4e**). These inversions may underlie phenotype or disease susceptibility differences, and further functional investigation is warranted to clarify their biological impact.

Using a pangenome graph built from 272 assemblies (including 8 Tibetan individuals), we identified 42,869 variable number tandem repeats (VNTRs), and 21,507 were located within 10 kbp of protein-coding gene bodies. Among these, 347 exhibited Tibetan-stratified frequency patterns (Wilcoxon rank-sum test, *p* < 0.05; TR disparity score (TRDS), TRDS ≥ 5), indicating substantial population-stratified variation for VNTRs (Figure 4f**; Supplementary Table 15**).

An example is a VNTR locus on chromosome 5 (chr5:1,324,770–1,326,443; 38 bp motif: GAAGGCACTGGGTGGGGGGCCGGGAGGGGCAGGGCGGG), located within the Intron 8 of *SLC6A3*, a dopamine transporter gene involved in synaptic signaling (Apsley et al., 2023). This VNTR displays a significantly elevated copy number in Tibetans compared to other populations (**Figure 4g-4i**), while it has been reported to affect gene expression and to associate with cocaine dependence in a Brazilian cohort (Guindalini et al., 2006). The mechanism driving this population-stratified pattern remains unknown, and further studies will be required to clarify whether it reflects high-attitude adaptation.

### Structural variants associated with high-altitude adaptation

Our Tibetan pangenome provide a unique opportunity to explore SVs underlying high-altitude adaptation. By constructing a combined pangenome from 70 Tibetans haploid assemblies and 204 Han Chinese haploids assemblies from the APGp1 dataset (Dongya Wu et al.), we identified 224 Tibetan-enriched SVs (TESVs), defined as SVs with a Tibetan allele frequency at least 0.2 higher than that in Han Chinese (**Fig. 5a; Supplementary Table 16; Methods**).

**Figure 5.**
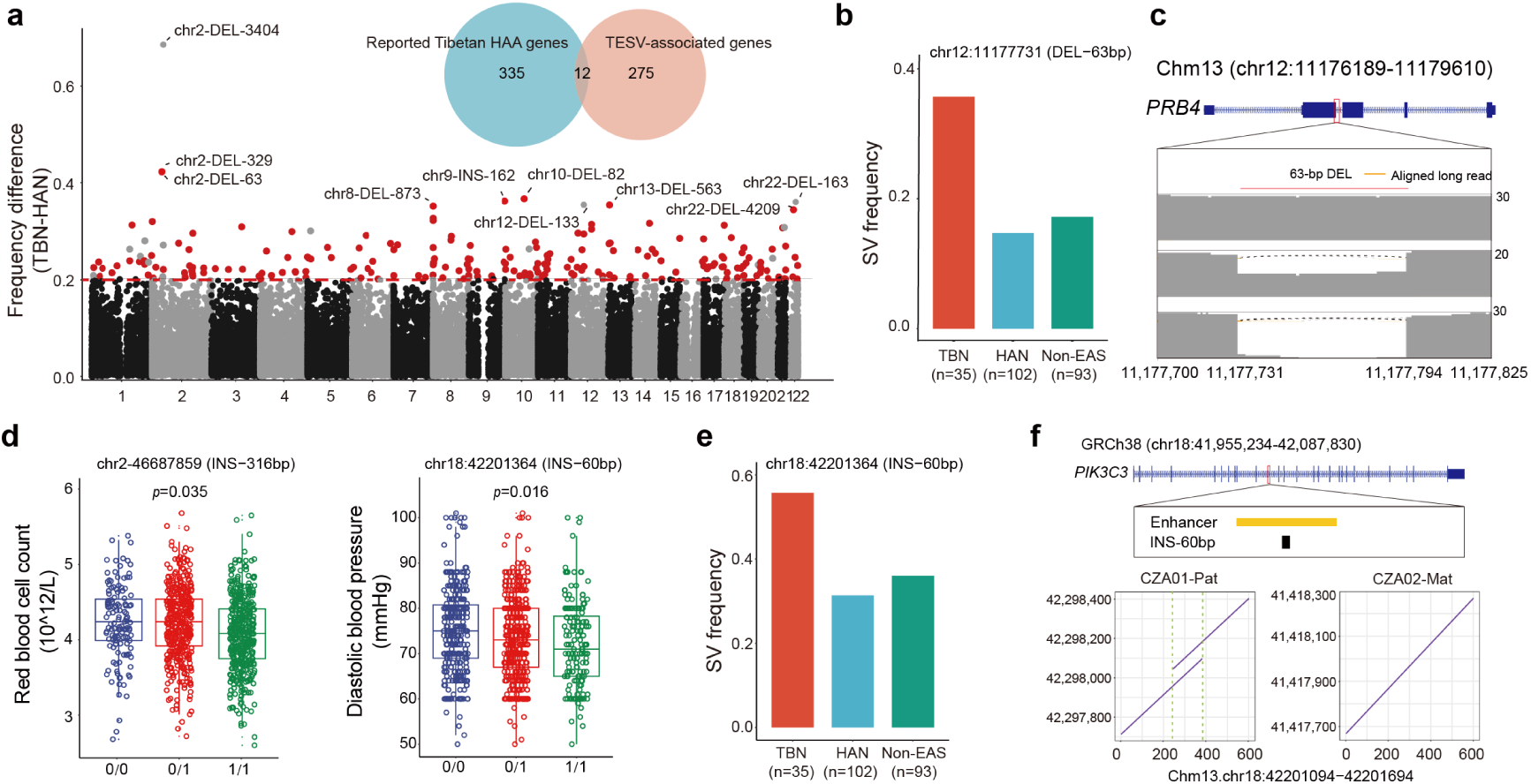
Adaptive SVs in Tibetan genomes. **a**, Genome-wide frequency differences of structural variants (SVs) between Tibetan (TBN) and Han Chinese (HAN) populations. The top ten signals are labeled with SV information (chromosome, SV type, and SV length in base pairs), with novel signals highlighted in red. The Venn diagram shows the overlap between previously reported SNV-based genes implicated in Tibetan high-altitude adaptation and the TESV-associated genes identified in this study. DEL, deletion; INS, insertion. **b,** Comparison of frequencies of the 63-bp deletion in the coding region of *PRB4* across populations. Non-EAS, non-East Asians. **c,** Genomic location and read-depth evidence of the 63-bp deletion within the *PRB4* gene. **d,** Significant associations under an additive genetic model were identified for red blood cell counts (left) and diastolic blood pressure (right) in Tibetans with the 316-bp insertion on chromosome 2 and the 60-bp insertion on chromosome 18, respectively. Genotype: 0/0 (non-SV), 0/1 (heterozygous SV), and 1/1 (homozygous SV). **e,** Comparison of frequencies of the 60-bp insertion on chromosome 18 between Tibetan and lowland populations (Han and non-East Asians, non-EAS). **f,** Genomic location (upper panel) of the 60-bp insertion on chromosome 18 and its flanking sequences in Tibetan populations. The insertion is overlapped with an annotated enhancer (based on ENCODE data). The dot-plot alignment (lower panel) highlights the 60-bp insertion by comparing two Tibetan assemblies with T2T-CHM13.

Among them, 17 TESVs were previously reported (He et al., 2020; Quan et al., 2021; Shi et al., 2023), including the well-known 3.4-kbp Tibetan-enriched deletion (TED) located ∼80 kbp downstream of *EPAS1*, which shows an allele frequency of 0.70 in Tibetans and 0.01 in Han Chinese (Lou et al., 2015).The remaining 207 TESVs represents newly identified TESVs, including 141 deletions and 66 insertions (**Supplementary Table 16**).

Genes located within 5-kbp downstream or upstream of TESVs were defined as TESV-associated genes (**Methods**). Totally, we identified 287 TESV-associated genes (**Supplementary Table 16**). Gene Ontology enrichment analysis revealed significant enrichment in three functional categories: glutamatergic synapse (FDR = 0.001), plasma membrane (FDR = 0.006), and GABAergic synapse (FDR = 0.011) (**Supplementary Table 17**), implying possible selection on the nervous system in Tibetans that was not proposed in previous SNV-based studies (Zheng et al., 2023).

To assess their relevance to hypoxic responses, we compared the TESV-associated genes with a curated set of 473 hypoxia-related genes (Peng et al., 2017). Ten genes overlapped with the 287 TESV-associated genes (odds ratio = 2.05, *p* = 0.03, Chi-squared test) (**Supplementary Fig. 10**), indicating a significant association with hypoxic regulation.

We further compared the TESV-associated genes to 347 genes previously reported showing strong signals of Darwinian positive selection in Tibetans (Zheng et al., 2023). Notably, 275 TESV-associated genes have not been reported before (**Fig. 5a**), suggesting that these SV-associated genes represent previously unrecognized candidates for high-altitude adaptation that could not be detected using SNV-based approaches. Among these TESVs, we identified a 63-bp deletion overlapped with the coding region of *PRB4* (Basic salivary proline-rich protein 4), which shows a markedly elevated frequency in Tibetans (0.36) compared with Han Chinese (0.15) and other global populations (0.17) (**Fig. 5b,5c**).

Utilizing the high-precision TESV breakpoint definitions from our pangenome, we performed SV genotyping in 1,001 Tibetans with short-read WGS data (Zheng et al., 2023), as well as 163 non-Tibetans with short-read WGS data from 1KGP dataset (Byrska-Bishop et al., 2022). In total, 222 TESVs were successfully genotyped and passed quality control (**Methods**). Given that 88 quantitative traits were phenotyped across these individuals, we conducted association analyses between the 222 TESVs and the quantitative traits. After Bonferroni correction, five TESVs showed significant associations with traits of heart and blood systems, supporting their roles in high-altitude adaptation (**Supplementary Table 18**):

- A 60-bp insertion (allele frequency: 0.56 in Tibetans; 0.31 in Han Chinese) located within an enhancer of the *PIK3C3* gene was significantly associated with lower diastolic blood pressure in Tibetans (effect size β = –0.17, Bonferroni-corrected *p* = 0.016) (**Fig. 5d–f**).
- A 758-bp deletion (frequency: 0.44 in Tibetans; 0.24 in Han Chinese) located ∼2 kbp from *GP5* and *ATP13A3* was significantly associated with lower platelet distribution width (β = –0.46, *p* = 0.016).
- A 55-bp deletion (frequency: 0.49 in Tibetans; 0.27 in Han Chinese) within the RNA gene *BDNF-AS* was significantly associated with increased interventricular septum thickness (β = 0.21, *p* = 0.017).
- A 316-bp insertion (frequency: 0.60 in Tibetans; 0.33 in Han Chinese) within the RNA gene *LOC124906001* showed significant association with lower red blood cell count (β = –0.09, *p* = 0.035) (**Fig. 5d**).
- A 309-bp insertion (frequency: 0.57 in Tibetans; 0.35 in Han Chinese) within the RNA gene *LOC105376496* was significantly associated with higher lymphocyte count (β = 0.09, *p* = 0.043).

### Archaic ancestry in complete genomes of Tibetans

To investigate archaic introgression in Tibetans, we applied SPrime (Browning et al., 2018) to autosome variants from the Tibetan pangenome based on 70 Tibetan haploid assemblies. In total, we detected 2.45 Gbp genome-wide archaic sequences, corresponding to 2,204 putative archaic-introgressed segments distributed broadly across the Tibetan genome (**Supplementary Fig. 11**). Across individuals, SPrime-inferred segments show an average of 70.1 Mbp of introgressed sequences per Tibetan genome, including 41.94 Mbp matching Neanderthal ancestry, 1.06 Mbp matching Denisovan ancestry, and 25.1 Mbp unknown ancestry (**Fig. 6a; Supplementary Table 19**).

**Figure 6.**
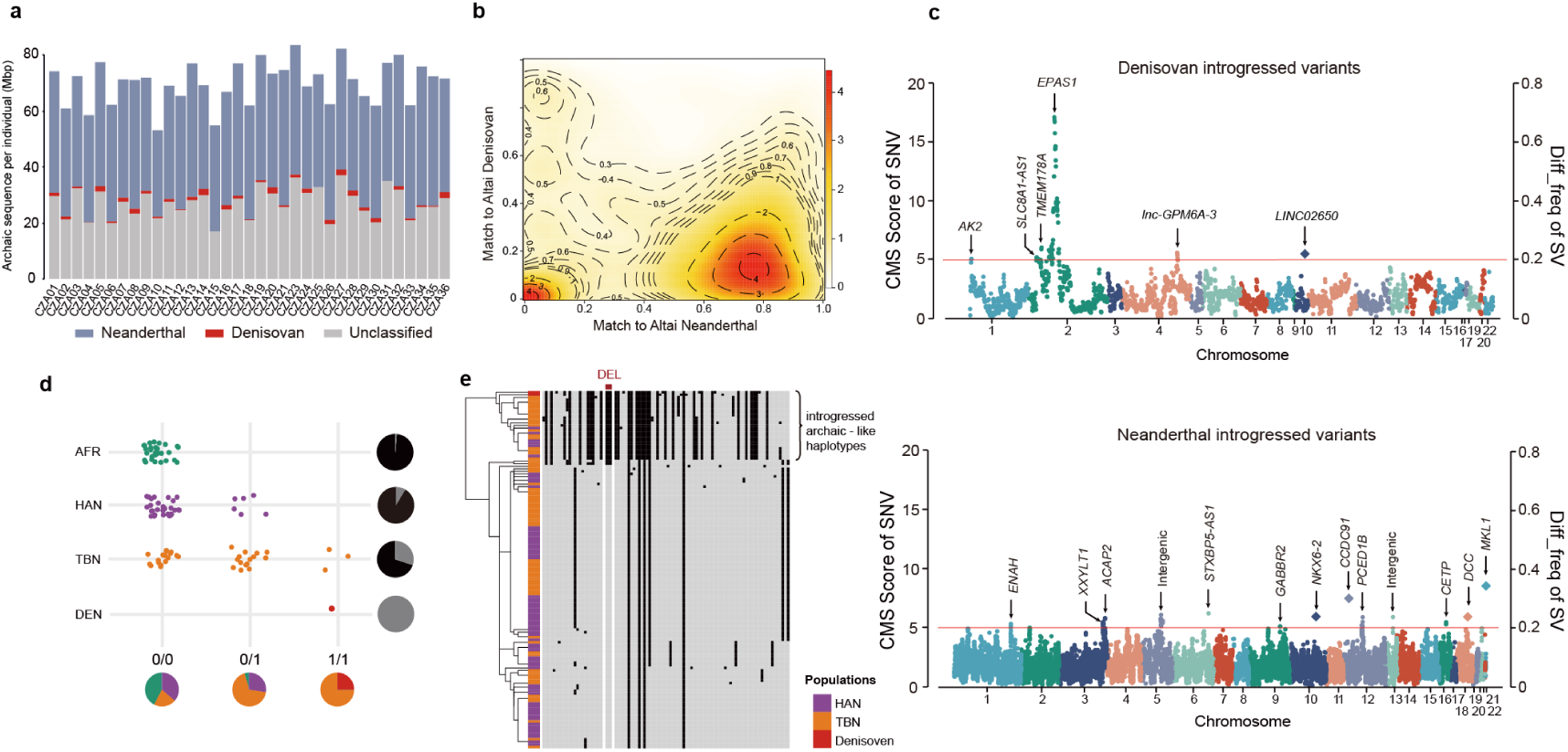
Landscape of archaic introgression in Tibetan genomes. **a**, Average amounts of the introgressed sequences per Tibetan individual, categorized by their affinity to the Altai Neanderthal and the Altai Denisovan genomes. **b,** Contour density plots showing joint match proportions of introgressed segments to the Altai Neanderthal and the Altai Denisovan genomes. Numbers within contour lines indicate the corresponding density levels. One Neanderthal (∼0.8 match rate), and two Denisovan (∼0.5 and ∼0.8 match rate) introgression events were identified in Tibetans. **c,** Signatures of positive selection in Tibetans for Denisovan-derived introgressed (upper panel) or Neanderthal-derived introgressed (lower panel) SNVs inferred by SPrime, evaluated using CMS (composite of multiple signals) scores. The red line denotes to the high CMS cutoff (CMS = 5). **d,** Genotyping results for the chromosome 10 Denisovan-introgressed deletion across global populations. Pie charts along the *x*-axis denote the population composition for each SV genotype (colors correspond to populations), whereas pie charts along the *y*-axis show deletion *versus* non-deletion allele frequencies within each population (gray and black represent deletion and non-deletion alleles, respectively). AFR, African; TBN, Tibetan; HAN, Han Chinese; NEA, Neanderthal; DEN, Denisovan. **e,** Hierarchical clustering of haplotypes spanning the adaptive-Denisovan-introgressed region on chromosome 10. Each row represents an individual haplotype. Columns correspond to the genotypes of the Tibetan-enriched 1,191-bp deletion, with gray and black denoting ancestral and derived alleles, respectively.

To infer the numbers of introgression events, we examined the joint distribution of match rates to the Altai Neanderthal and Altai Denisovan genomes for putative archaic segments (Browning et al., 2018). We observed a major cluster with a high Neanderthal match rate (∼0.8), and two distinct clusters with Denisovan match rates (∼0.5 and ∼0.8) (**Fig. 6b**). These patterns support one pulse of Neanderthal introgression and two pulses of Denisovan introgression in Tibetans, consistent with previous findings of more than one Denisovan introgression in eastern Asia based on short-read sequencing of Tibetan (Zhang et al., 2021), East Asian (Browning et al., 2018) and Southeast Asian genomes (He et al., 2025).

To look for the signatures of “borrowed” fitness due to archaic introgression in Tibetans, we checked whether variants located within the introgressed segments matching Neanderthal-or Denisovan-derived alleles showed evidence of strong positive selection (CMS score >5) and enriched in Tibetans (**Methods**). In total, we identified 19 candidate adaptive introgressed regions, including 6 Denisovan-derived regions and 13 Neanderthal-derived regions (**Fig. 6c; Supplementary Table 20, 21**). One marked example is a ∼651 kbp Denisovan-introgressed region on chromosome 2, containing *EPAS1, TMEM247, ATP6V1E2, RHOQ, PIGF* and *CRIPT* (**Fig. 6c; Supplementary Table 20**). This region has been repeatedly reported as a classic instance of adaptive introgression in Tibetans (Huerta-Sánchez et al., 2014; Shi et al., 2023; Zhang et al., 2021; Zheng et al., 2023). Importantly, we identified 18 additional novel introgressed regions showing modest but detectable signatures of positive selection, including 5 Denisovan-introgressed and 13 Neanderthal-introgressed (**Figure 6c; Supplementary Figure 12; Supplementary Table 20, 21**). These introgression regions overlapped with genes associated with diverse phenotypes, including lung function, blood pressure, bone density, and nervous system development.

Notably, we found five introgressed regions harbor five TESVs. For example, we identified a 1,191 bp deletion as Denisovan introgressed SV in Tibetans. This SV showing a high frequency divergence between Tibetans and Han Chinese (0.30 *versus* 0.09) (**Fig. 6d**). Our hierarchical clustering and network analysis confirmed the tight clustering of this haplotype with the Denisovan sequence (**Fig. 6e**), supporting the proposed adaptive introgression in Tibetans. Besides, we identified four Neanderthal-introgressed SVs enriched in Tibetan populations (**Supplementary Table 21**). Among them, we identified a 163-bp deletion in *MKL1*—showing an allele frequency difference of 0.227 between Tibetans and Han Chinese—which exhibits clear Neanderthal introgression signatures (**Supplementary Table 21**). This deletion was proven in our previous study that significantly associated with lower systolic pulmonary arterial pressure, one of the key adaptive physiological traits in Tibetans (He et al., 2020). These observations suggest that introgressed SVs might be contribute to hypoxia adaptation in Tibetans.

## Discussion

Our study provides the most complete and comprehensive genomic portrait to date of an indigenous high-altitude human population, demonstrating how telomere-to-telomere assembly and pangenome frameworks can transform our understanding of human adaptation. By generating 35 fully phased Tibetan genomes, including eight that reach near-T2T completeness, and integrating them into a population-specific pangenome graph, we characterize the full spectrum of genetic diversity in Tibetans, including sequences previously inaccessible in standard reference genomes. These resources illuminate how complex genomic regions, SVs, and archaic introgression have shaped the evolutionary trajectory of high-altitude populations.

A major advance of this work is the resolution of genomic regions that have long resisted complete assembly. We show that Tibetan telomeres exhibit considerable structural heterogeneity, dominated by canonical repeat tracts but also featuring noncanonical “blocky” motifs present across global populations. Tibetan centromeres reveal both expected stability within individuals and notable Tibetan-specific HOR configurations, including a disrupted α-satellite array on chromosome 5 and a distinctive HOR structure on chromosome Y. These features likely arose from population-specific mutational dynamics or drift in repetitive DNA, and their biological consequences remain to be explored. The eight fully assembled Tibetan Y chromosomes further exposes substantial structural variability, including recurrent inversions at known hotspots, including a 4.2-Mbp inversion enriched in the Tibetan-specific haplogroup D-M533. These results underscore the functional and evolutionary richness of highly repetitive regions, which until recently were largely omitted from studies of human adaptation.

By constructing a Tibetan-specific pangenome graph, we uncover over 120 Mbp of non-reference sequences, representing previously unsolved core regions in the Tibetan population. These sequences reveal marked population stratification between Tibetans and Han Chinese and highlight complex regions—such as the dynamic β-defensin gene cluster at 8p23.1, where Tibetan genomes harbor extensive structural diversity. The overlap between Tibetan-specific pangenome bubbles and archaic haplotypes further suggests that archaic introgression affect genomic complexity in high-altitude populations. These insights reinforce the value of population-tailored pangenomes for capturing biologically important variation that is systematically missed by linear references.

A central goal of this study was to evaluate the contribution of SVs to high-altitude adaptation. By integrating our pangenome-defined SV catalog with population allele frequencies, archived whole-genome data, and extensive population phenotyping, we identify 224 TESVs, the majority of which are new. These TESVs implicate new biological pathways in Tibetan adaptation, including synaptic and neuro-related processes, functional domains that have not emerged prominently from SNV-based scans of selection. Although the mechanistic links between neuronal signaling and hypoxic adaptation are not well understood, accumulating evidence supports a role for hypoxia in modulating neurovascular coupling, ventilatory responses, and cardiorespiratory integration (Ojikutu et al., 2025; Terraneo et al., 2017), raising the possibility that structural variation influences neural components of high-altitude physiology.

Our genotype–phenotype analyses in 1,164 Tibetans and Han further identify five TESVs associated with hematological and cardiovascular traits central to high-altitude adaptation. These include variants near *PIK3C3, GP5, ATP13A3*, and noncoding RNAs influencing diastolic blood pressure, platelet morphology, interventricular septum thickness, and red blood cell count. Although effect sizes are modest, they are consistent with polygenic architectures and demonstrate that SVs contribute functionally relevant variation beyond canonical SNVs. The discovery of TESV-associated genes not captured in previous positive-selection scans emphasizes the necessity of integrating SVs into evolutionary analyses and raises the prospect that additional adaptive loci remain to be uncovered as long-read datasets expand across global populations.

Our archaic introgression analysis reveals that Denisovan- and Neanderthal-derived sequences are widespread in Tibetan genomes and extend far beyond the well-known *EPAS1* locus. The joint match-rate patterns support a single Neanderthal pulse and two Denisovan introgression events (Browning et al., 2018), now resolved with complete haplotypes. We identify 14 candidate regions of adaptive introgression, including previously unreported Denisovan tracts in genes related to hematopoiesis, cardiovascular function, and hypoxia responses. Notably, several Tibetan-enriched SVs also carry archaic signatures, including a Denisovan-derived 1.2-kbp deletion and multiple Neanderthal-derived SVs linked to cardiopulmonary and neurodevelopmental pathways. These findings indicate that archaic introgression contributed multi-scale genetic variants, both SNVs and SVs, that were subsequently shaped by high-altitude selection in Tibetans.

Despite the transformative scope of these data, several limitations warrant caution. First, although our near-T2T assemblies provide deep insight into the Tibetan genome, additional high-coverage assemblies from diverse Tibetan subpopulations will further resolve region-specific variation shaped by varied environmental conditions and distinct demographic histories. Second, while our association analyses identify trait-relevant SVs, experimental assays, including reporter constructs, genome editing, and molecular phenotyping under hypoxic conditions, will be essential to establish causal mechanisms. Third, although our pangenome captures extensive non-reference sequence content, highly repetitive regions (e.g., rDNA arrays and pericentromeric satellite blocks) remain difficult to fully resolve and integrate into the graph. As a result, variants within these regions are largely missing from our pangenome representation, underscoring the need for further advances in assembly and graph-construction algorithms.

In sum, our study demonstrates the power of combining T2T assemblies with a population-specific pangenome to reveal previously concealed variation and reinterpret the genetic basis of human adaptation. This work lays a foundational resource for high-altitude genomics and provides a generalizable blueprint for constructing pangenomes in other environmentally or culturally unique human populations. As T2T and pangenome technologies continue to advance, we anticipate a rapid expansion in our understanding of how complex genomic variation, repetitive DNA, and archaic inheritance shape human diversity and evolutionary resilience.

## Acknowledgements

We thank all participants in this project. This study was supported by National Key R&D Program of China (2025YFC3410300 to B.S. and Y.H.), National Natural Science Foundation of China (32288101 to B.S.; 32522022, U25A20648 and 32170632 to Y.H.), Major Scientific Project of Yunnan Province (202305AH340007 to B.S.), Yunnan Revitalization Talent Support Program Innovation Team (202405AS350008), Yunnan Scientist Workshops (to B.S.), Science and Technology General Program of Yunnan Province (202301AW070010 to Y.H.).

## Author contributions

B.S., Y.H., and Y.M. conceived and designed the study. B.S., Y.H., and Y.M. coordinated and supervised the project. Y.H., X.Q., C.C., Ouzhuluobu and Y.Z. contributed to sample collection, personnel coordination and volunteer organization in the collection work. Y.H., K.L. prepared the samples and processed to sequencing. K.L., J.L., D.W. and G.Z. contributed to data processing, quality control, variant analysis, genome assembly. K.L. and L.M. contributed to pangenome construction and analyses. L.M. contributed to analysis of telomere, centromere, and chromosome Y. K.L., L.M., J.S. and Y.H. contributed to structural variants analysis. F.Z., L.F., J.H. and Y.M. contributed to complex variants analysis. J.S. contributed to archaic introgression analysis. B.S. and Y.H. were responsible for ethical, legal and social implications. Y.H. and B.S. wrote the manuscript. K.L., L.M., F.Z., Y.M., Y.L. and L.C. edited the manuscript. All authors read and approved the final manuscript.

## Additional information

Supplementary information is available for this paper at Supplementary Information.

## Data availability

All data generated in this study are deposited in public repositories hosted by the National Genomics Data Center. Whole-genome sequencing reads are available in the Genome Sequence Archive for Human (GSA-Human: https://ngdc.cncb.ac.cn/gsa-human/), genome assemblies in the Genome Warehouse (GWH: https://ngdc.cncb.ac.cn/gwh/), and identified variants in the Genome Variation Map (GVM: https://ngdc.cncb.ac.cn/gvm/), under BioProject accession PRJCA043737. These datasets include short-read WGS data, PacBio HiFi reads, ultra-long ONT reads from 35 trios (105 individuals), as well as the resulting genome assemblies and pangenome graph.

In accordance with international standards for protecting participant privacy, raw sequencing data and individual-level genome data are accessible through controlled access for non-profit scientific research. Researchers may request access by submitting an application, including a research proposal and institutional IRB approval, to the Data Access Committee of the Kunming Institute of Zoology, Chinese Academy of Sciences (KIZ, CAS).

## Methods

### Data production

A total of 35Tibetan trios (105 individuals) were recruited from a hospital in Lhasa, Xizang Autonomous Region, China (elevation: 3,650 m). All procedures involving human participants were reviewed and approved by the Institutional Review Board of the Kunming Institute of Zoology, Chinese Academy of Sciences (Approval ID: KIZRKX-2025-002). High-molecular-weight genomic DNA was extracted from peripheral blood, and three sequencing platforms were used to generate PacBio HiFi, Oxford Nanopore ultra-long, and PCR-free short-read data. All experimental procedures adhered to the ethical standards of the Responsible Committee on Human Experimentation.

#### HiFi sequencing

Peripheral blood samples were thawed at 37 °C, and high-molecular-weight genomic DNA was extracted using the Qiagen MagAttract HMW DNA kit following the manufacturer’s instructions. DNA quantity was measured using a Qubit fluorometer. The DNA was sheared to a target size of 15–18 kbp using g-TUBEs, followed by end repair, A-tailing, and ligation of SMRTbell adapters using the SMRTbell Prep Kit 3.0. Libraries were purified and assessed for quality and fragment size using an Agilent 2100 Bioanalyzer. Sequencing was performed on PacBio Revio platforms to generate circular consensus sequencing (CCS) HiFi reads.

#### Ultra-long ONT sequencing

Genomic DNA was extracted from peripheral blood mononuclear cells (PBMCs) isolated from whole blood using the NEB Monarch Tissue DNA Extraction Kit. DNA quality and concentration were assessed by Qubit fluorometry and electrophoresis. Ultra-long read libraries were prepared using the Oxford Nanopore SQK-LSK114 kit following a modified hexammine-cobalt–based NEMO bead protocol, including DNA repair, adapter ligation, and bead-based purification. Final libraries were loaded onto primed PromethION R10.4.1 flow cells and sequenced on the Oxford Nanopore PromethION platform with extended runtime to maximize ultra-long read yield.

#### PCR-free NGS sequencing

Genomic DNA was extracted using a magnetic bead-based automated protocol (CWBio CW2361S). For PCR-free library construction, 50–1000 ng of high-quality genomic DNA was enzymatically fragmented and size-selected using magnetic beads to generate insert sizes of 450–600 bp (high input) or 600–750 bp (low input). Fragmented DNA underwent end repair, A-tailing, and ligation with MGIEasy PF adapters. Libraries were purified, denatured, and circularized; residual linear DNA was digested with exonuclease to produce single-stranded circular (ssCir) libraries. Library quantity was measured with a Qubit ssDNA assay, and size distribution was assessed on an Agilent Bioanalyzer. Libraries with ≥75 fmol yield were sequenced on the MGI DNBSEQ-T7 platform.

#### Genome assembly and assessment

Before generating the assemblies, we detected and removed reads containing PacBio adapters using a bash script from the HiFiAdapterFilt repository and using NanoFilt (De Coster et al., 2018) to retain ONT reads above 100 kbp for genome assembly to enhance genome continuity. The command was as follows:

*bash hifiadapterfilt.sh -l 44 -m 97 -t 64 -p sample.hifi*

*gunzip-c sample.ont.fq.gz | NanoFilt -l 100000 | gzip > sample.ont.100k.fq.gz*

We carried out diploid assembly using Hifiasm (0.24.0-r702) (Cheng et al., 2024) trio binning module when parental short reads are available, and obtained 54 haplotype-resolved assemblies for the 27 samples. Command as follows:

*yak count -k31 -b37 -t16 -o pat.yak paternal.fq.gz yak count -k31 -b37 -t16 -o mat.yak maternal.fq.gz*

*hifiasm -o sample.asm -t 32 -1 pat.yak -2 mat.yak --ul sample.ont.100k.fq.gz sample.hifi.filt.fq.gz*

Hifiasm produces assemblies in GFA format. Each diploid GFA file was then converted to FASTA format using the following command:

*for pre in ‘ls | grep “p_ctg.gfa” | perl -npe “s/.p_ctg.gfa//”’; do awk ’/^S/ (print “>”$2“\n”$3)’ pre.p_ctg.gfa | fold > pre_ctg.fa;done*

We then carried out assembly polishing using nextpolish2 (v0.2.0) (Hu et al., 2024) which takes a genome assembly file, a HiFi mapping file and one or more k-mer dataset files from short reads as input and generates the polished genome. The command was as follows:

*nextPolish2 -t 16 sample.hifi.filt.mapped.bam sample.asm sample.k21.yak sample.k31.yak > sample.np2.fa*.

We next applied Inspector (Chen et al., 2021) to assesses the misassemblies before and after polish to determine whether the assembly polishing improved the assembly quality. The assessment command of assembly errors using Inspector was as follows:

*python inspector.py -c sample.asm.fa -r sample.hifi.filt.fq.gz -o sample/ --datatype hifi -t 64*.

Assembly QV was determined using k-mer-based tools. Yak’s QV estimation applies separately on each haplotype. The k-mer databases for Yak (Cheng et al., 2021) were generated using the following command:

*yak count -t32 -k21 -b37 -o sample.yak < (zcat sample.ngs.fq.gz) < (zcat sample.ngs.fq.gz)*.

QV estimation using Yak was generated with the following command:

*yak qv -t 32 -p sample.yak sample.np2.fa > sample.asm.yak.qv.txt*.

We also applied QUAST (Gurevich et al., 2013) to assess the contiguity, completeness and duplication ratio for each polished assembly. The assessment command of QUAST was as follows:

*python quast.py sample.np2.fa -r reference.fa -o sample/ --large --est-ref-size 3100000000 --no-icarus -t 64*.

Switch errors and hamming errors were evaluated for maternal and paternal haploid assembly by using yak *trioeval*. We used Flagger (v0.3.2) (Liao et al., 2023), a read-alignment-based pipeline, to detect putative assembly errors in the diploid assemblies. Flagger utilizes the alignments of long reads to the diploid assembly, fits the mapping coverage distribution by constructing mixture models, and classify genomic blocks into different categories reflecting assembly accuracy at those locations, including correct regions (haploid) and regions that may contain errors.

### Genome annotation

#### Telomere identification and classification

Telomeric regions were annotated using the VGP telomere pipeline (https://github.com/VGP/vgp-assembly/tree/master/pipeline/telomere) with the parameters 0.4 15000. Briefly, telomeric repeats were detected in 1-kbp windows (200-bp sliding step) within the terminal 15 kbp of each chromosome. Windows containing ≥10 % of the canonical hexamer “TTAGGG” were retained. Putative telomeric intervals were extracted with BEDTools (v2.31.1) (Quinlan and Hall, 2010). Sequence identity heatmaps were generated with ModDotPlot (v0.9.4) (Sweeten et al., 2024) (parameters: *Static Mode, --kmer 21, --identity 86, --delta 0.5, -r* 1000).

Three telomere architectures were observed: i) C-type: Continuous TTAGGG arrays; ii) F-type: Fragmented arrays with short, interspersed variants disrupting the canonical repeat; iii) B-type: Blocks dominated by non-canonical motifs. B-type telomeres were operationally defined by two criteria: a) non-TTAGGG/non-complementary motifs constituted ≥5 % of the telomeric sequence. b) total length ≥2 kbp to ensure structural completeness. To objectively separate C-type and F-type telomeres, three unsupervised methods were applied to the sequence-identity distributions: i) Kernel density estimation (KDE) with a Gaussian kernel; the optimal density threshold was 0.40; ii) K-means clustering with the elbow criterion; empirical thresholds were 0.7255 (Tibetans, n = 40) and 0.7249 (APGp1 samples, n = 160); iii) Gaussian mixture models (GMM) assuming two latent distributions; the derived cutoffs were 0.7584 (Tibetan dataset) and 0.7513 (APGp1 dataset). Owing to non-normal distributions in telomere length across several chromosomes, inter-chromosomal differences were evaluated with the Kruskal–Wallis test. When a significant omnibus result was obtained (*p* < 0.05), pairwise comparisons were carried out using Dunn’s post-hoc test with Bonferroni correction to identify specific chromosome pairs exhibiting significant differences. All statistical analyses were performed in R (v4.4.1).

#### Centromeric repeat annotation and ROH diversity analysis

Centromeric repeats were firstly annotated with RepeatMasker (v4.1.6) (https://repeatmasker.org/). Regions identified as “SAR” were annotated as HSat1A, and regions annotated as “HSATI” were annotated as HSat1B, and regions annotated as “GSAT” were annotated asγSat, and regions annotated as “BSR” were annotated as β-Sat.HSat2 and HSat3 arrays were independently identified and classified with the custom script Assembly_HSat2and3v2.pl (Altemose et al., 2022).

α-satellite monomers were annotated with the centromere annotation pipeline (https://github.com/ssssyq1123/centromere_annotation). All previously identified centromeric sequences were filtered to retain only α-satellite monomers. After re-formatting the coordinate files with a Python script (mon2stv.py), monomers located within ROH were subjected to k-mer classification using stv.sh (https://github.com/fedorrik/stv). Subsequently, additional satellite classes residing in the same ROH intervals—βSat, γSat, HSat1A, HSat1B and HSat2—were appended to the dataset. Summary statistics and visualizations were generated with custom R scripts. Identical procedures were applied to the publicly available centromere annotations of T2T-CHM13 (v2.0) and 160 APGp1 individuals. The resulting ROH satellite profiles were compared to characterize Tibetan-specific centromeric diversity patterns. For the test of the length difference of ROH in different chromosomes, the Kruskal-Wallis test and the Dunn test were used.

#### Annotation of Y chromosomes

The eight Tibetan Y chromosome assemblies were annotated by transferring features from T2T-CHM13 (v2.0) (Nurk et al., 2022) (https://github.com/marbl/CHM13). Amplicon, Pseudo-autosomal Region (PAR), sequence-class, important-region and inverted-repeat BED files were downloaded from the T2T-Y and mapped to each Tibetan assembly with liftoff v1.6.3 (default parameters). Pairwise collinearity between each Tibetan chrY and the T2T-Y reference was compared with syri (v1.6.3) (Goel et al., 2019). Chromosomes were first aligned with minimap2 (v2.28) (Li, 2018) to produce SAM files, which were then processed by syri under default settings. Structural differences were visualized with plotsr (v1.1.1) (Goel and Schneeberger, 2022). Global sequence-identity heatmaps were generated using ModDotPlot (v0.9.4) in Static Mode (*--kmer 21 --identity 86 --delta 0.5 -r* 1000).

Single-nucleotide variants (SNVs) were called with DeepVariant (v1.6.1) (Yun et al., 2021), and the T2T-Y as reference genome. Complex regions (telomeres, centromeres, heterochromatic blocks and PARs) were excluded with BEDTools (v2.31.1) (Quinlan and Hall, 2010). The resulting VCFs were converted to GRCh37 with GATK (v4.6.1.0) (Van der Auwera and O’Connor, 2020) (LiftoverVcf). Haplogroups were annotation with yhaplo (v2.1.14) (https://github.com/23andMe/yhaplo) (--anc_snps). A subset of high-quality SNVs located in the same non-repetitive regions across all samples was extracted. vcf2phylip was used to generate a PHYLIP alignment, and the optimal evolutionary model was selected automatically with IQTree2 (v2.3.6) (Minh et al., 2020) (-m MFP). A maximum-likelihood phylogeny was inferred under the best-fit model with 1,000 ultrafast bootstrap replicates.

### Tibetan pangenome graph

#### Pangenome construction

We constructed pangenome variation graphs for 35 Tibetan individuals using the Minigraph-Cactus pipeline (Hickey et al., 2024), which integrates structural variants (SVs) and single-nucleotide variants (SNVs) into a unified graph representation. Both GRCh38 and T2T-CHM13 were included as reference genomes, with T2T-CHM13 serving as the primary backbone. Graph construction followed the procedures described in the Minigraph–Cactus documentation.

#### Pangenome growth curve

Pangenome growth was assessed using Panacus (v0.2.3) (Parmigiani et al., 2024). We quantified the accumulation of genomic content across samples, defining >95% presence as core, >5% as common, and the presence of a single haplotype as singleton. The following command was used:

*panacus ordered-histgrowth -c bp -t 32 -l 1,2,4,67 -H -e TBN35-v2-mc-chm13.new.Pline.txt TBN35-v2-mc-chm13.gfa.gz*

#### Graph-based small variant calling

Variant sites within the Minigraph–Cactus graphs were identified using vg deconstruct (Garrison et al., 2018) (v1.56.0). Alternative alleles at multiallelic SV sites were grouped by length, and variant sites were classified into: single nucleotide variants (SNV), small insertion-deletions (INDEL, <50bp and ≥1bp), multiple nucleotide polymorphisms (MNP), and structural variants (SV ≥50bp) sites. Multiallelic sites were subsequently decomposed into biallelic records using: *bcftools norm -m -any*.

#### Bubble counts statistics

We constructed a Minigraph-Cactus pangenome graph incorporating all 70 Tibetan assemblies and 204 Han Chinese assemblies from APGp1, using T2T-CHM13 (v2.0) as the reference backbone. From the resulting pangenome variant VCF file, we extracted a subset of 35 Tibetan individuals and a randomly selected subset of 35 Han individuals for comparative analyses. Using VCFtools, we performed a sliding-window scan on both populations with a window size of 1 Mbp and a step size of 100 kbp. For each window, we counted the number of bubbles and calculated population difference in bubble counts as:

*d_(TBN-HAN)_ = Tibetan bubble counts – Han bubble counts*

We selected the top 0.1% of windows with the highest *d_(TBN-HAN)_* values and compared them with the genome-wide distribution using a Wilcoxon test. This analysis yielded 27 regions with significantly elevated bubble counts in Tibetans relative to Han.

To refine the spatial resolution of these signals, we conducted a second sliding-window analysis using a 100-kbp window with a 10-kbp step. We recalculated *d_(TBN-HAN)_* for each window and focused on the 1-Mbp candidate regions identified in the first scan. After removing redundancies due to overlapping windows, we obtained 21 non-overlapping 10-kbp regions that consistently showed higher bubble counts in Tibetans.

### SV discovery

#### Pangenome graph-based SV calling

We used vg deconstruct to identify variant sites in the Minigraph–Cactus graph. Large spurious deletions were removed using **vcfbub** with the parameters: “*vcfbub -l 0 -r 100000*”. To generate non-redundant SV callset, we used *truvari* to collapse similar SVs using the command: *truvari collapse -r 500 -p 0 -P 0.5 -s 50 -S 100000 - k common -i MC.vcf.gz -c collapsed.MC.vcf.gz -o MC.collapse.vcf.gz*.

#### Read mapping-based SV calling

SVs were also identified directly from PacBio HiFi reads using two independent pipelines. Reads were aligned to the T2T-CHM13v2 reference genome using Minimap2 (v2.2) for Sniffles2 (Smolka et al., 2024) and pbmm2 (https://github.com/PacificBiosciences/pbmm2) for pbsv, both with default parameters. SV calling was then performed using Sniffles2 (v2.2) and pbsv, requiring each SV to: i) be supported by ≥10 HiFi reads; ii) be ≥50 bp and ≤1 Mb in size, and iii) contain precise breakpoint annotations. To avoid false positives in highly repetitive regions, we removed all SVs located within centromeres, telomeres, and the surrounding ±5 Mbp flanking regions.

#### Assembly-based SV calling

We additionally identified SVs directly from the haplotype-resolved assemblies using two assembly-based callers. Phased Assembly Variant Caller (PAV) (v2.3.4) (Ebert et al., 2021) and svim-asm (Heller and Vingron, 2021) were used to detect simple SVs—including insertions, deletions, and inversions—by aligning each assembly to the T2T-CHM13 (v2.0) reference genome. To minimize artifacts from highly repetitive regions, we removed all SVs located within centromeres, telomeres, and the surrounding ±5 Mbp regions. We retained variants ranging from ≥50 bp to ≤1 Mbp for downstream analyses.

#### Variants novelty evaluation

To assess the novelty of small variants and SVs in our Tibetan call set, we compared the 35-individual Tibetan variant set against publicly available variant catalogs from HGSVC3 (Logsdon et al., 2025) and HPRC (Liao et al., 2023). For SNVs, we required exact positional and allelic matches. For SVs and INDELs, we used *bedtools intersect* with the parameters *-f 0.5 -r* to identify reciprocal overlaps. Two SVs (or INDELs) were considered the same variant only if their reciprocal overlap exceeded 50% of the length of each variant.

#### Identification of Tibetan-enriched SVs

The pangenome graph provided high-confidence SVs with precise breakpoint resolution, enabling robust SV genotyping using deep whole-genome sequencing data and facilitating frequency comparisons between Tibetan and Han Chinese populations. We constructed a Minigraph-Cactus pangenome using 35 Tibetan and 102 Han Chinese samples and screened for SVs showing a Tibetan–Han allele frequency difference greater than 0.2. All candidate SVs were subjected to rigorous manual validation. Breakpoints and genotypes were examined by aligning each sample’s PacBio HiFi reads to the T2T-CHM13 (v2.0) reference genome. SVs that were not supported by read-mapping evidence were further evaluated using assembly-based visualization with mummer4 (Marçais et al., 2018). Through this combined approach, we identified 224 high-confidence Tibetan-enriched SVs (TESVs), which were retained for downstream phenotype association analyses.

#### Association analysis between SV and phenotype

Using the high-precision TESV breakpoints derived from the pangenome, we performed SV genotyping on 1,001 previously published Tibetan genomes sequenced with short-read whole-genome data using Pangenie (Ebler et al., 2022). In total, 222 TESVs from 953 individuals were successfully genotyped and passed quality control. The quality control filters included a missing rate < 0.05, Hardy–Weinberg equilibrium *P* < 1×10⁻¹⁰, and minor allele frequency > 0.01. We analyzed 88 quantitative traits measured across all individuals, removed outliers exceeding 3 standard deviations, and assessed associations between SVs and phenotypes using plink *--linear*, incorporating the top 10 principal components, with altitude and age as covariates.

#### Assessment of gene copy number variation

We employed Liftoff (v1.6.3) to estimate gene copy number for each assembly, utilizing the GENCODE (v38) database for annotation. The parameter *“-sc 0.95”* was applied to ensure a minimum sequence similarity threshold of 95% for identified gene copies. The execution command was as follows:

*liftoff -p 32 -sc 0.95 -copies -db GENCODE_V38.db -u <sample.unmapped> -o <sample.gff3> -polish <sample.fa> GRCh38.fa*

In the resulting GFF3 output files for each assembly, the *“extra_copy_number”* field (representing any value greater than 0) indicates the corresponding gene copy number variation.

The initial results underwent stringent filtering, retaining only exonic and protein-coding genes. Alignments meeting thresholds of ≥95% sequence similarity and ≥90% of the original gene length were considered candidate copy number genes. Furthermore, we performed manual inspection using IGV to exclude potential false-positive calls generated by Liftoff.

### Inversion calling

#### Generation of inversion callset

We applied several complementary genomic structure variants caller to construct independent inversion callsets using PAV (long-read-based phased assemblies) (Ebert et al., 2021), PBSV (https://github.com/PacificBiosciences/pbsv) and LSGvar (https://github.com/Hanjunmin/LSGvar). To create a final nonredundant inversion callset outside of centromeric regions, we merged inversion calls based on different technologies using bedtools (Quinlan and Hall, 2010) with following options: *bedtools intersect -a callset1.bed -b callset2.bed -f 0.5 -F 0.5 -wa -wb*. Merging of overlapping inversion calls was done in the following priority order: PAV, PBSV and LSGvar.

#### Inversion filtering and refinement of the breakpoints

We refined inversion breakpoints with dot plots. We first extracted inversion region in the T2T-CHM13 reference and used minimap2 (Li, 2018) to identify the corresponding region in each assembled haplotype. We expanded 100-kbp upstream and downstream of the inversion breakpoints in reference and corresponding region in each haplotype. *Nucmer* was used to create pairwise sequence alignments between these extracted regions and their reference counterpart in T2T-CHM13. Alignments were finally visualized as dot plots using mummerplot (Marçais et al., 2018). Manual curation of inversion breakpoints was performed using breakpoints identified in these alignments, with dot plots and saffire for visual guidance.

#### Population-stratified inversion

We intersected the Tibetan inversion callset with the APGp1 inversion dataset. An inversion was considered orthogonally supported if it exhibited ≥50% reciprocal overlap with an inversion reported using an orthogonal genomic technology in a public dataset. We further identified Tibetan-enriched inversions using a two-tailed Fisher’s exact test. In total, this analysis revealed 23 Tibetan-enriched inversions and 5 Tibetan-specific inversions (Supplementary Table 18).

#### Inversion site verification

To validate large inversion calls, we firstly performed alignments of long reads (Supplementary Fig. 11). Taking the ∼449kb inversion on 7q11.21 as an example, high-quality alignments of phased ONT reads from maternal and paternal haplotypes on T2T-CHM13 were visualized in the IGV (v2.19.1), displaying collapse signals around the breakpoints, as an indicative of inversion.

### VNTR identification

To identify variable number tandem repeats (VNTRs), we used a Pan-VNTR pipeline from (Han et al.) to annotate motifs onto a pangenome graph constructed from APGp1 (Dongya Wu et al.), including 8 Tibetan individuals. VNTRs with motif ≥ 7bp, copy numbers > 2 and lengths ranging from 150bp to 10k were selected, resulting in 42,869 VNTRs loci. Of the identified VNTRs, 21,507 loci (50.2%) were associated with protein-coding genes.

To explore population-specific differences, we calculated copy number divergence (defined as a difference of > 5 copies) and calculated Tandem Repeat Disparity Score (TRDS) to identify significant VNTR loci between Tibetan and other superpopulations. After keeping loci where motif annotations covered greater than 80% of the total VNTR length, we identified 625 Tibetan-specific VNTR loci with statistically significant differences (*p* < 0.05) while 347 (55.5%) located in protein-coding genes.

### Archaic introgression detection

We investigated Neanderthal and Denisovan introgression in Tibetans using 35 nearly complete genomes. As a modern human outgroup, we used 108 Yoruba individuals (1000 Genomes Project). High-coverage Altai Neanderthal and Altai Denisovan reference genomes were obtained from https://www.eva.mpg.de/genetics/genome-projects/. Archaic introgressed alleles and the genomic segments defined by these alleles were identified using SPrime (https://github.com/browning-lab/sprime). The detection step was performed using the following command:

*java -jar sprime.jar gt=genotype.vcf.gz outgroup=outgroup_id.txt map=GRCh38.map out=out.file chrom=chr*

To generate individual-level introgression calls, we reconstructed the archaic ancestry status along each genome from the SPrime output. Consecutive archaic-specific alleles occurring on the same haplotype were merged into putative introgressed segments using:

*map_arch --kp --sep ’\t’ --tag archaic_name --mskbed mask.bed.gz --vcf archaicfile.vcf.gz --score scorefile > out.chr.mscore*

To reduce false positives, we required each segment to contain at least 30 archaic-allele sites. Neanderthal-derived segments were defined as those with a match rate >0.6 to the Altai Neanderthal genome and <0.4 to the Altai Denisovan genome. Denisovan-derived segments were defined as those with a match rate >0.4 to the Altai Denisovan and <0.3 to the Altai Neanderthal. Segments with >0.7 match rate to the Denisovan genome were classified as high-affinity Denisovan introgression.

#### Adaptive introgression

We identified putatively adaptively introgressed SNVs using the composite of multiple signals (CMS) approach. CMS scores for ∼9 million Tibetan SNVs were obtained from previous large-scale Tibetan WGS data (Zheng et al., 2023). CMS scores were then assigned to all archaic-introgressed SNVs identified by SPrime, including Neanderthal- and Denisovan-derived variants. SNVs with a CMS score >5 were considered candidate adaptive introgressed alleles. Results were visualized as Manhattan plots using R.

To detect adaptive introgressed structural variants (SVs), we genotyped 224 Tibetan-enriched SVs (TESVs) in four high-coverage archaic genomes—Altai Denisovan, Altai Neanderthal, Vindija Neanderthal, and Chagyrskaya Neanderthal—as well as in African outgroup samples (YRI from 1000 Genomes Project). Candidate introgressed SVs were defined as those with at least one archaic-matched allele observed in Tibetans but absent in the African outgroup.

---

**Supplementary Fig. 1.**
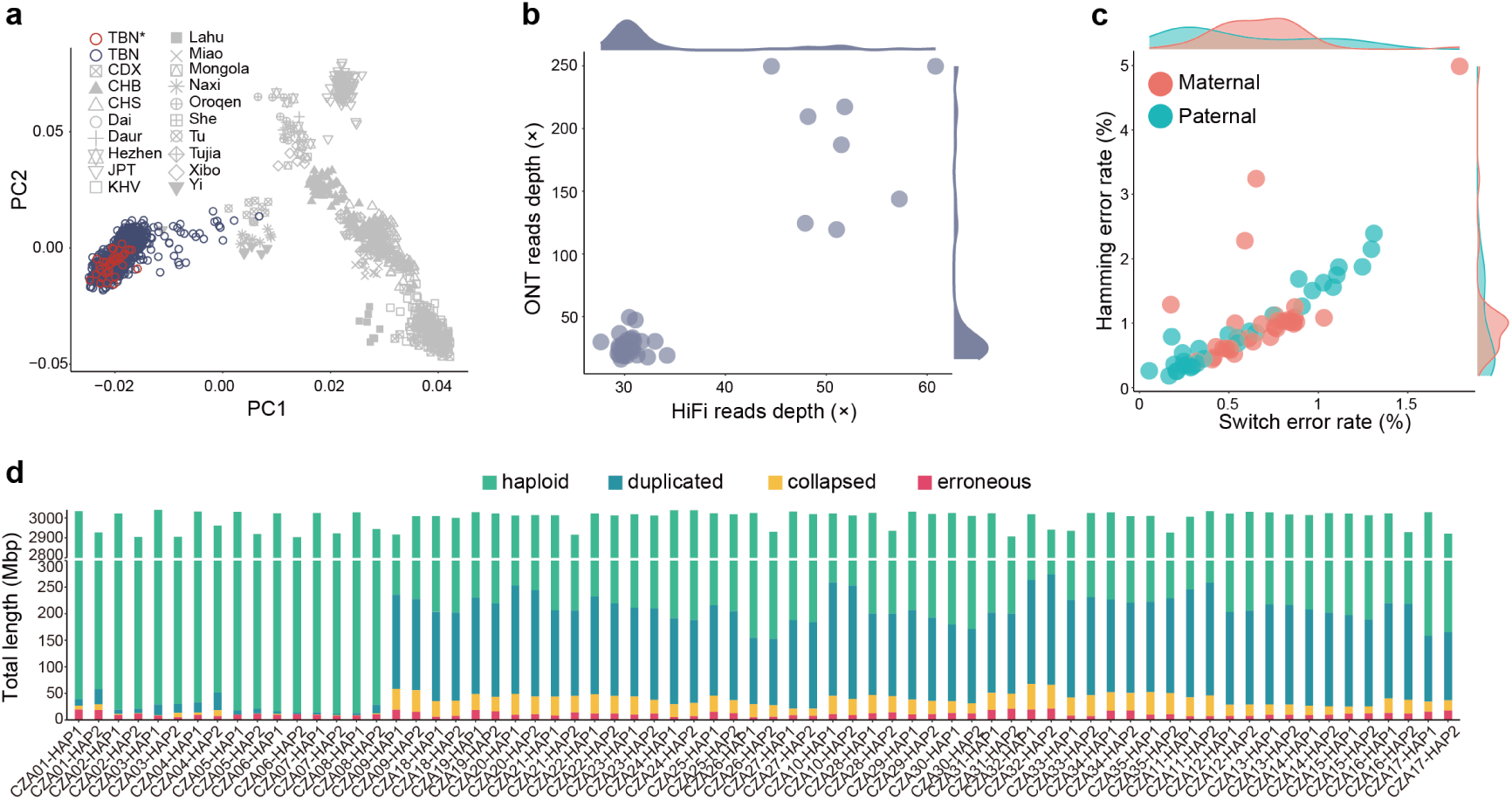
Sample information, sequencing data, and assembly quality for Tibetan genomes. **a**, Principal component analysis (PCA) of 105 Tibetan individuals (35 trios; red) shown alongside East Asian populations (gray) and 1,001 previously published Tibetans (blue; Zheng et al. 2023). **b**, Sequencing depth of PacBio HiFi and ultra-long ONT reads for the 35 Tibetan samples. **c**, Phasing accuracy estimated by Yak, showing switch error rates versus Hamming error rates. **d**, Assembly reliability assessed by read-mapping support. Green segments indicate haploid regions with high reliability. The y-axis is broken to highlight the predominance of reliable haploid blocks and the distribution of unreliable segments.

**Supplementary Fig. 2.**
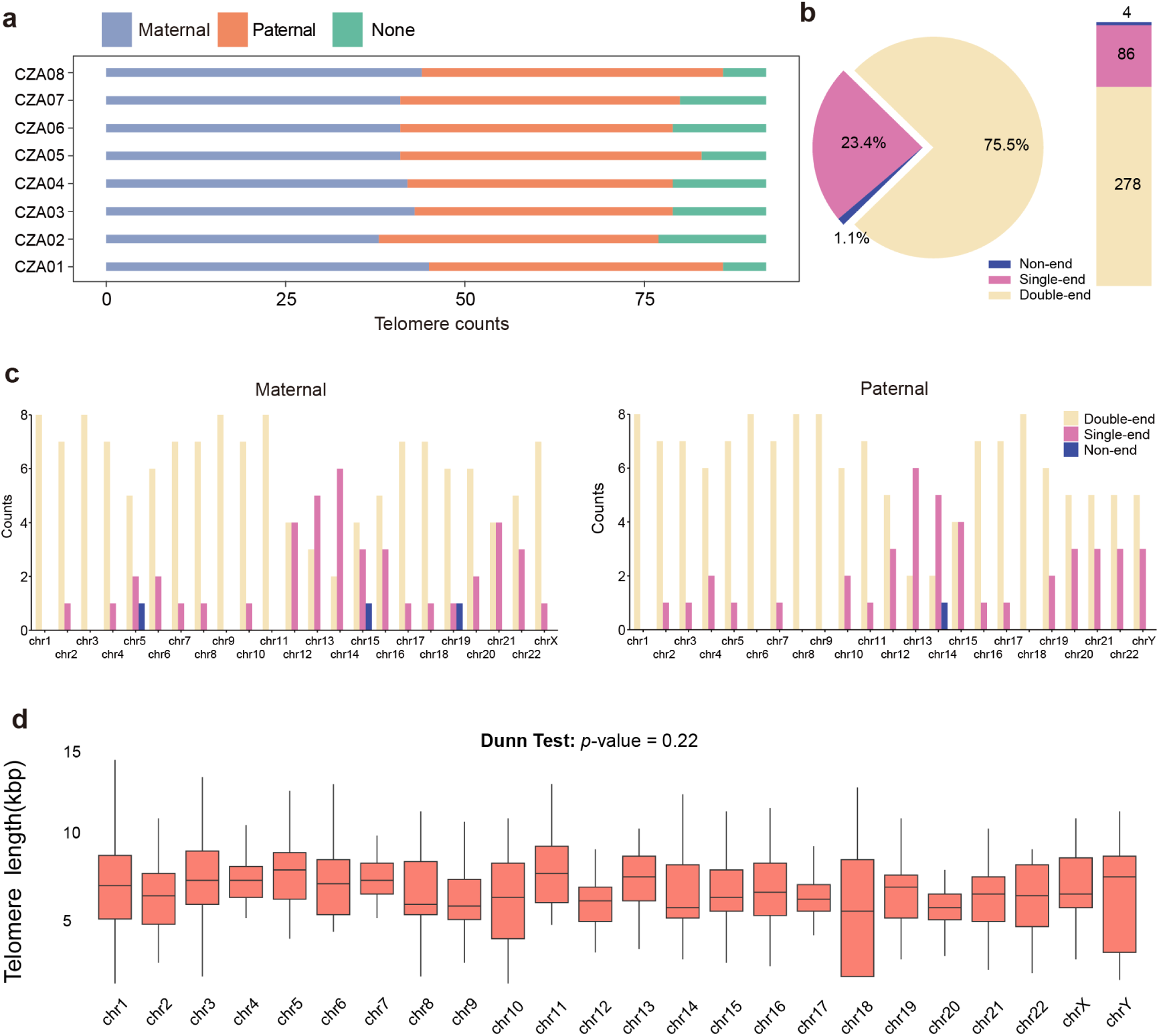
Telomere length distribution in Tibetans. **a**, Summary of telomere annotations for eight Tibetan genomes. Telomere counts for maternal and paternal haplotypes are shown, ‘None’ indicates the telomere unable to annotation. **b**, Proportion of telomere annotations per chromosome. Non-end indicates chromosomes without annotated telomeres at either end; single-end indicates annotation at only one end; and double-end indicates successful annotation at both chromosomal ends. **c**, Chromosome-level telomere annotation counts across the eight maternal assemblies and eight paternal assemblies, respectively. **d**, Inter-chromosomal telomere length differences assessed using the Kruskal–Wallis test, followed by Dunn’s post-hoc test with FDR correction.

**Supplementary Fig. 3.**
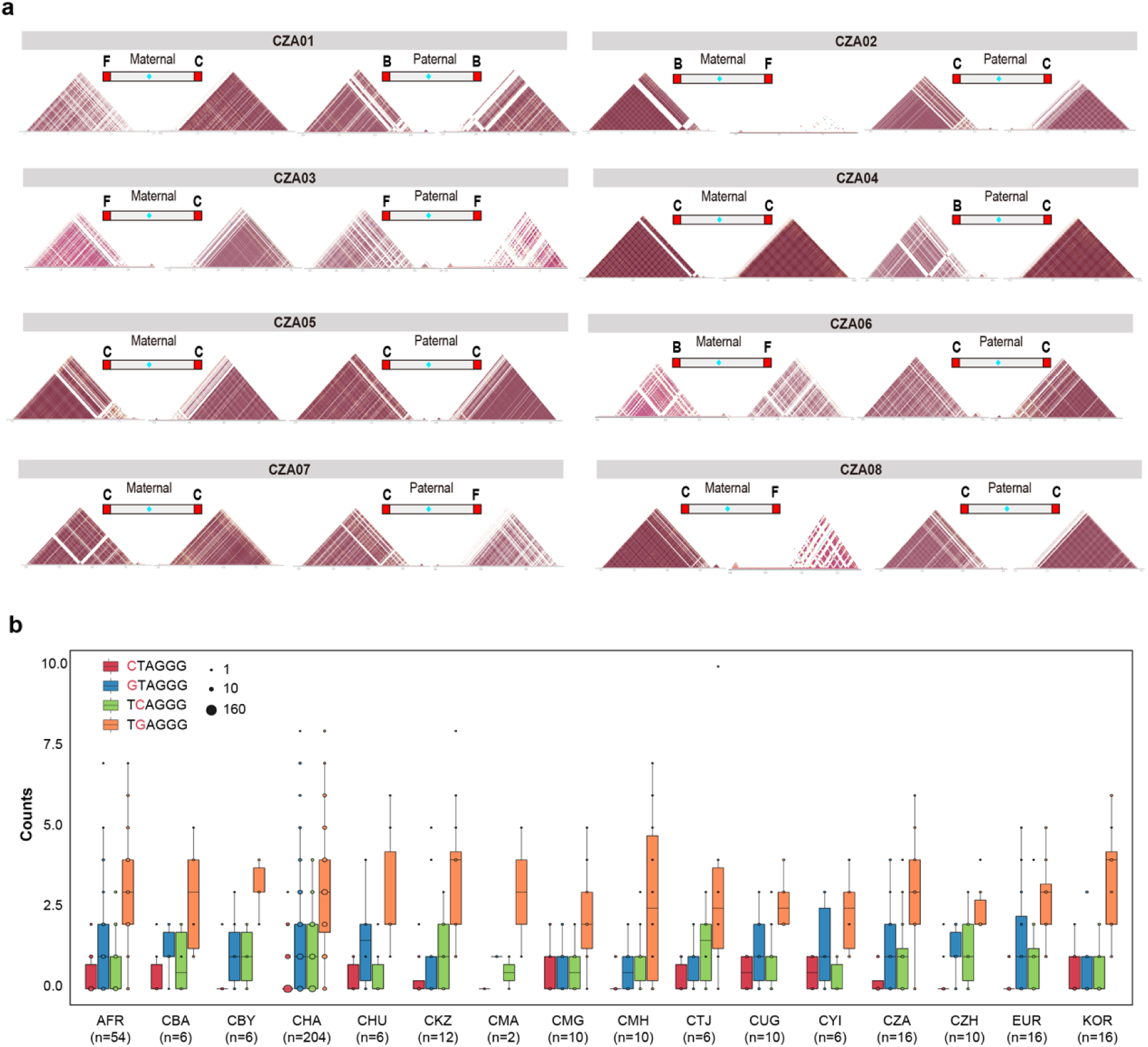
Sequence identity and global distribution of telomere subtype B. **a**, Heat map of sequence identity for telomere region in 16 Tibetan haplotype genomes (take chromosome 1 as example), with corresponding subtype classifications (C, B, F) indicated. **b**, Counts of subtype B type telomeres across worldwide populations (AFR, EUR from HPRC dataset; the other from APG dataset). CBA, Bai Chinese; CBY, Buyi Chinese; CHA, Han Chinese; CMA, Man Chinese; CMG, Mongol Chinese; CTJ, Tujia Chinese; CUG, Uygur Chinese; CYI, Yi Chinese; CZH, Zhuang Chinese; KOR, Korea; AFR, African; EUR, European.

**Supplementary Fig. 4.**
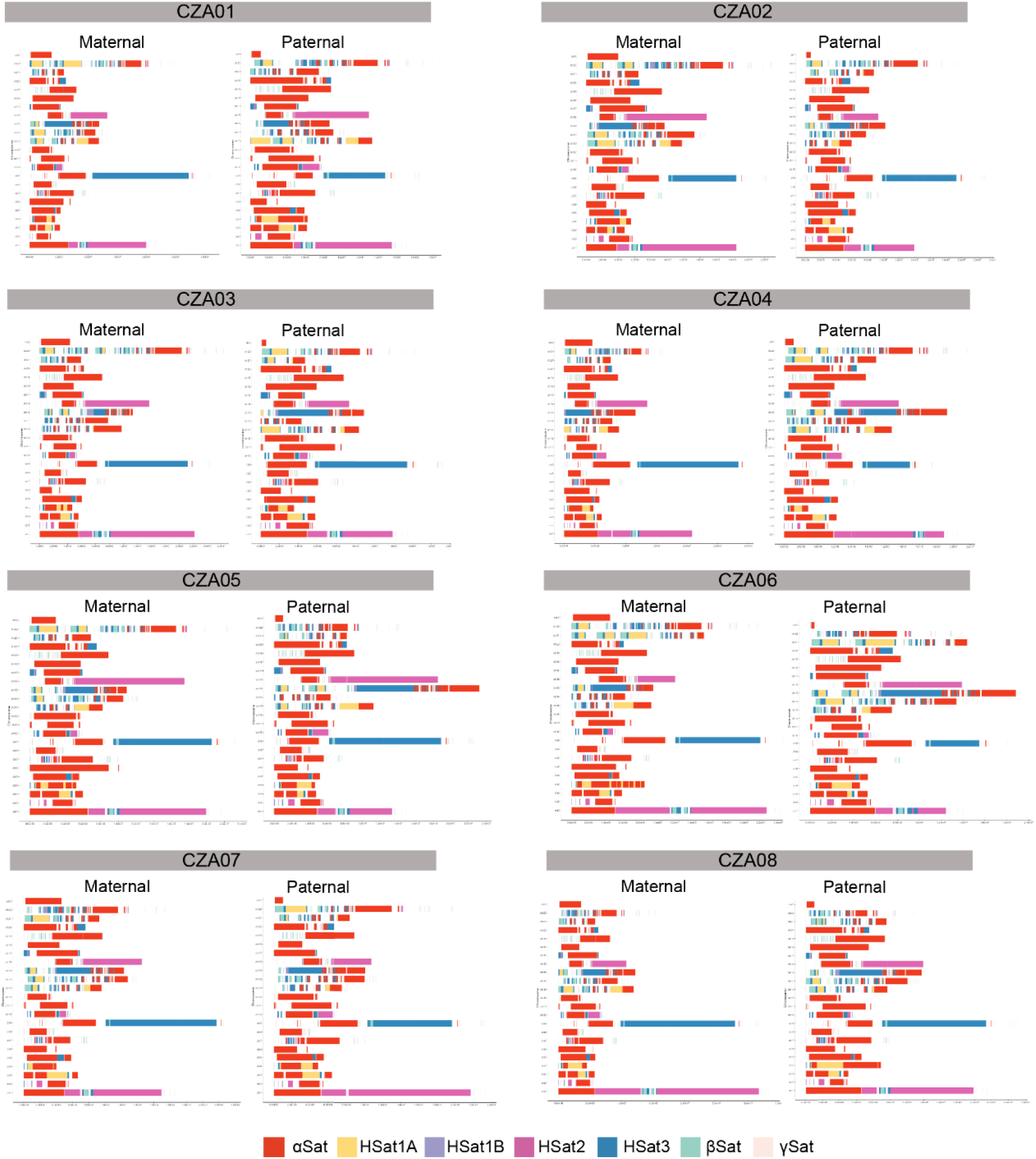
Color-coded map of peri/centromeric satellite DNA arrays across all Tibetan assemblies. The colored blocks represent different satellite arrays, including α-satellite (αSat), β-satellite (βSat), H-satellite1A (HSat1A), H-satellite1B (HSat1B), H-satellite2A (HSat2A), H-satellite3 (HSat3), and γ-satellite (γSat).

**Supplementary Fig. 5.**
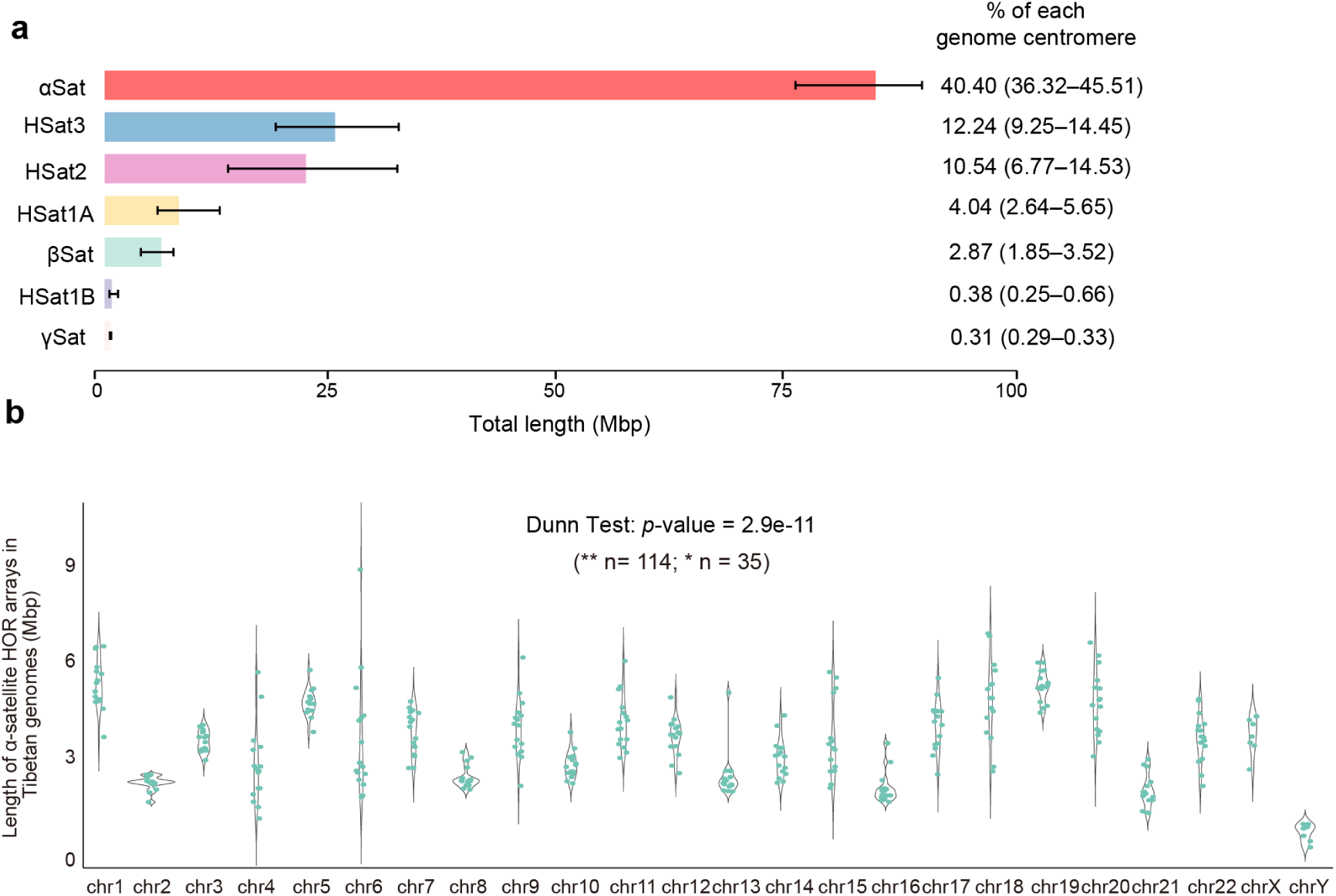
Centromere satellite-DNA landscape and *α*-satellite HOR organization in Tibetan genomes. **a**, Barplots of the total lengths of each major satellite family in eight Tibetan genomes. The proportion of each type in each whole centromere is shown in the barplot right. **b**, Length of α-satellite higher-order repeat (HOR) arrays for each complete and accurately assembled centromere in Tibetan haplotype genomes. Comparison across chromosomes was performed using Dunn’s test. 114 chromosome pairs exhibited extremely significant divergence (\*\**p* < 0.01) and 35 pairs showed significant divergence (\**p* < 0.05).

**Supplementary Fig. 6.**
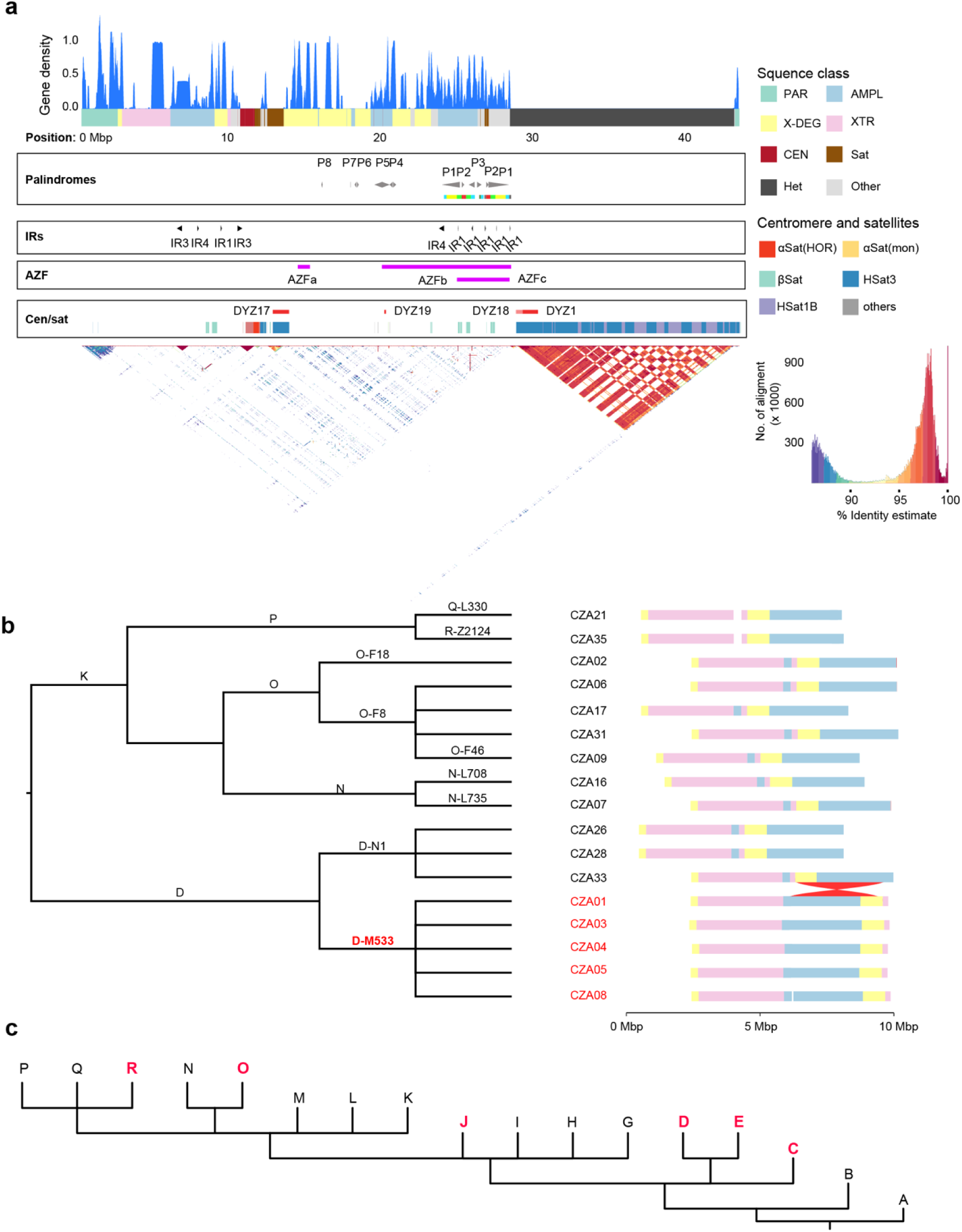
Structure and phylogeny of a complete Tibetan Y chromosome. **a**, Gene density plot of the euchromatic region of the Y chromosome, showing enriched protein-coding genes. Sequence classes, palindromes, inverted repeats (IRs), Azoospermia factor regions (AZFa, AZFb, AZFc), centromere, satellite repeats, and sequence identity are annotated. **b**, Phylogeny of 17 Tibetan Y-chromosomal haplogroups from 35 Tibetan individuals. A 4.2 Mbp inversion prevalent in the Tibetan D-M533 lineage is shown to the right of the phylogenetic tree. **c**, Composite Y-chromosome phylogeny incorporating all publicly available data. Haplogroups containing the 4.2 Mbp inversion are highlighted in red.

**Supplementary Fig. 7.**
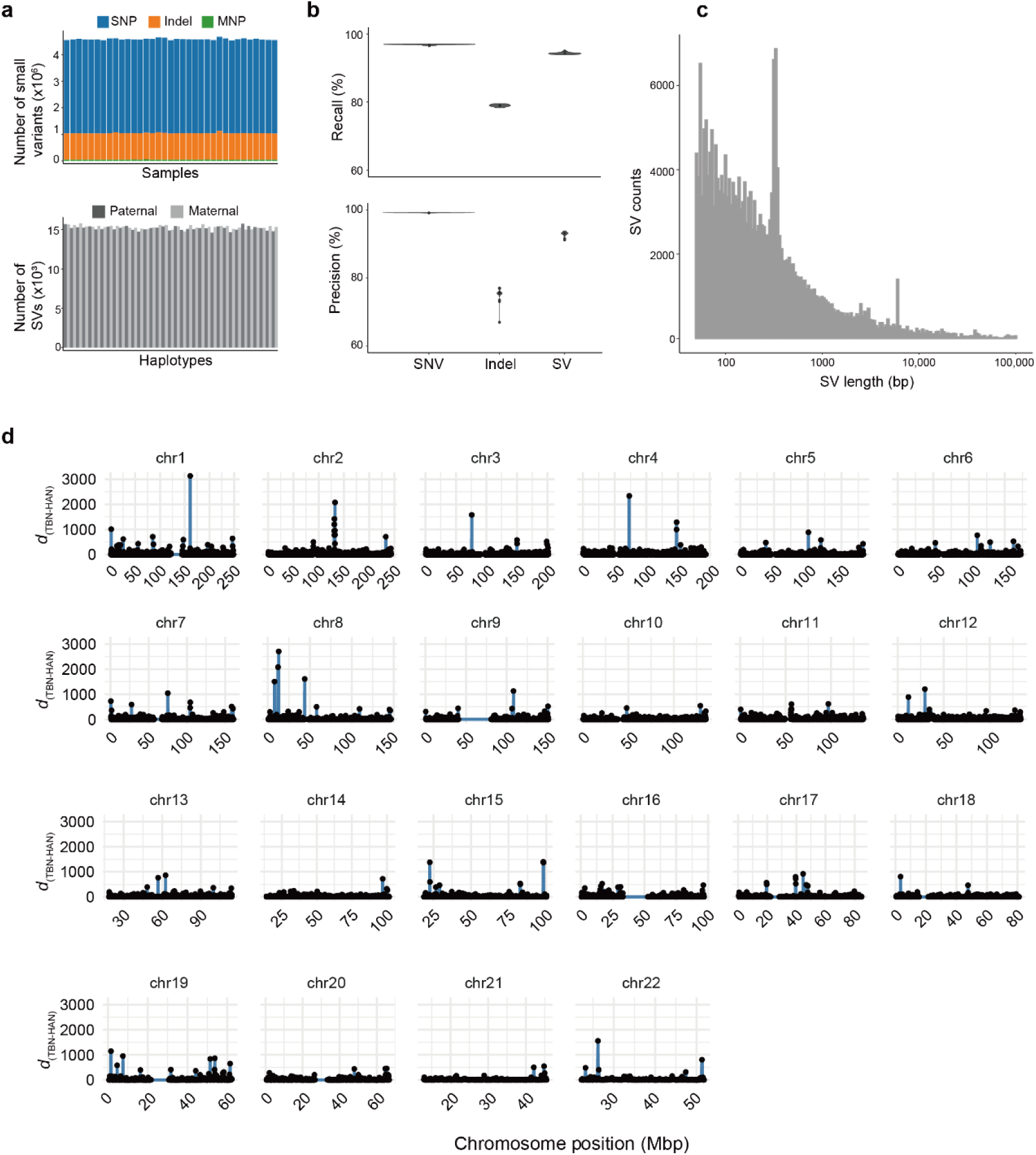
Statistics of the Tibetan pangenome. **a–b**, Numbers of small variant sites and structural variants (SVs) carried per individual. MNP, multinucleotide polymorphism. **b**, Precision and recall of autosomal small variants and SVs in the pangenome relative to consensus call sets. Small variants are benchmarked against HiFi–DeepVariant calls; SVs are benchmarked against the consensus of four SV callers (Methods). Analyses exclude centromeres, rDNA, telomeres, segmental duplications, and simple repeats. **c**, Length distribution of SVs in the graph-based pangenome reference. **d**, Genome-wide differences in bubble counts between Tibetans and Han Chinese (*d*_(TBN–HAN)_) in 1-Mb windows.

**Supplementary Fig. 8.**
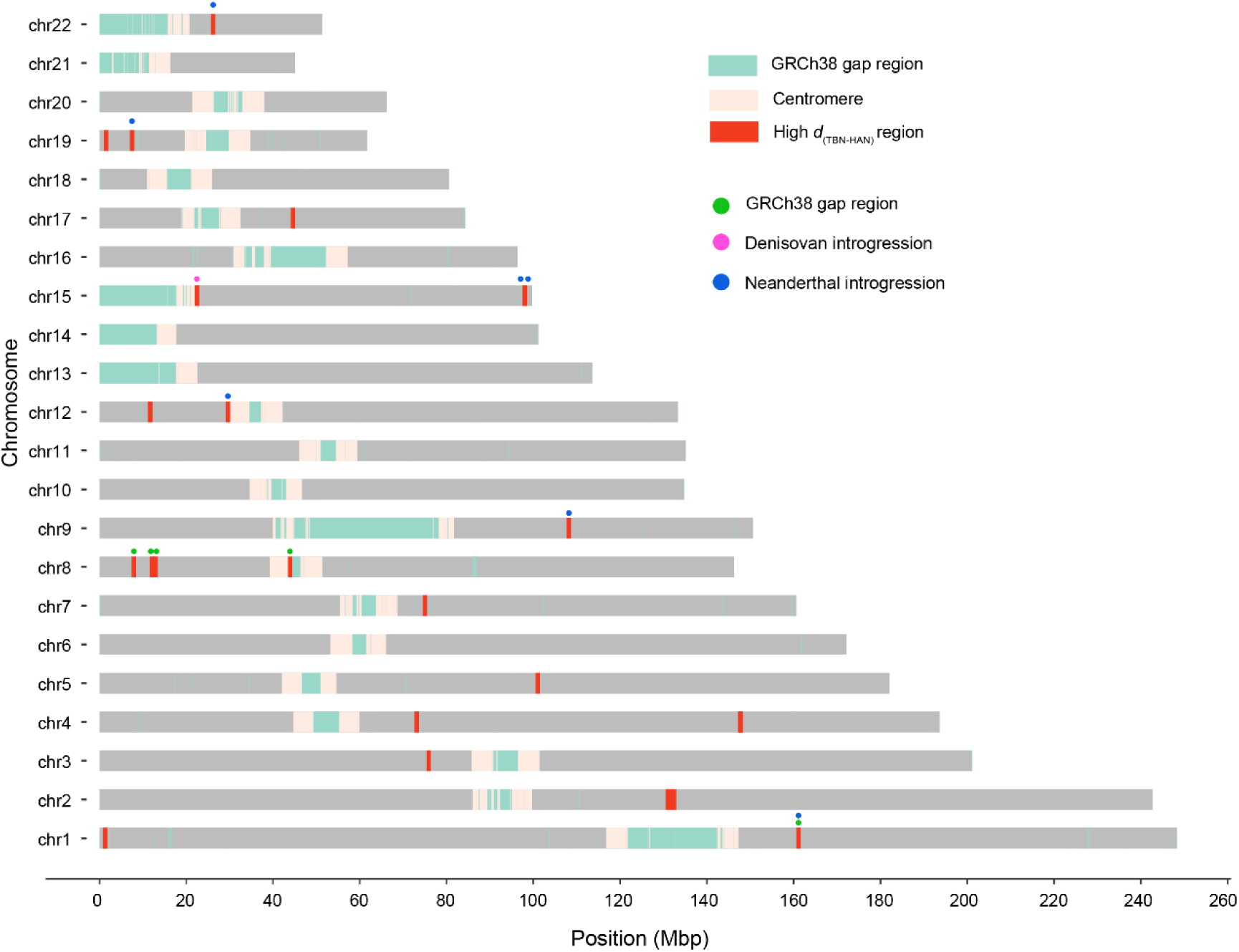
Distribution of population-stratified genomic regions based on bubble-count differences between the Tibetan and Han Chinese pangenomes. Red 1-Mbp windows denote regions where Tibetans show significantly higher bubble counts than Han Chinese. Green windows mark GRCh38 gaps or misassemblies that are resolved in the T2T-CHM13 reference. Blue dots indicate Tibetan-enriched bubble regions that coincide with these corrected GRCh38 gaps. Pink dots highlight regions that additionally overlap putative Denisovan introgression tracts, whereas green dots mark those intersecting Neanderthal-introgressed segments.

**Supplementary Fig. 9.**
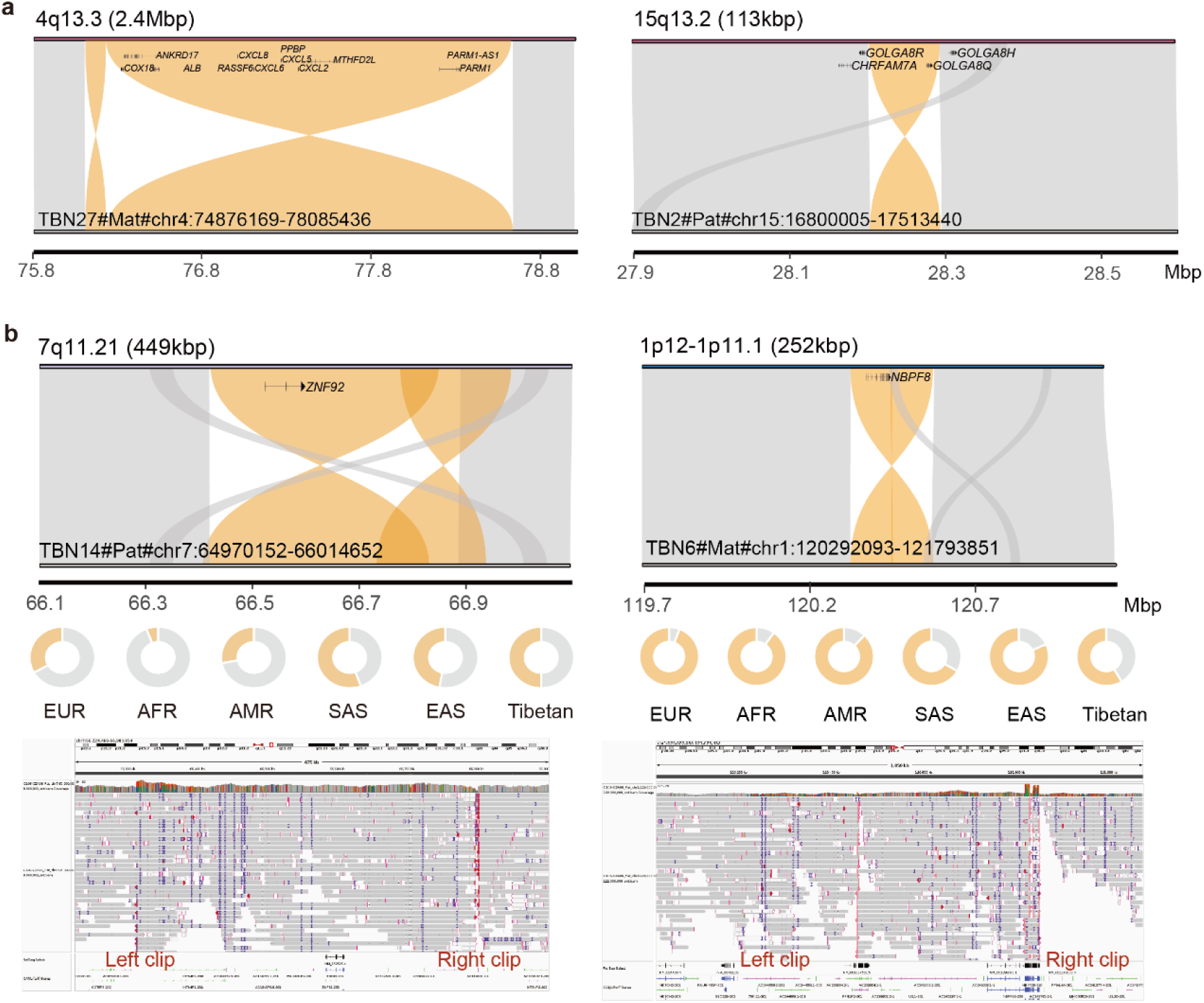
Validation of large inversions. **a**, Synteny plots of two Tibetan-specific inversions relative to the T2T-CHM13 reference. **b**, Large inversions commonly observed in Tibetans (>2 assemblies), validated by aligning phased ONT reads to T2T-CHM13. The genomic locations of the inversion breakpoints are highlighted.

**Supplementary Fig. 10.**
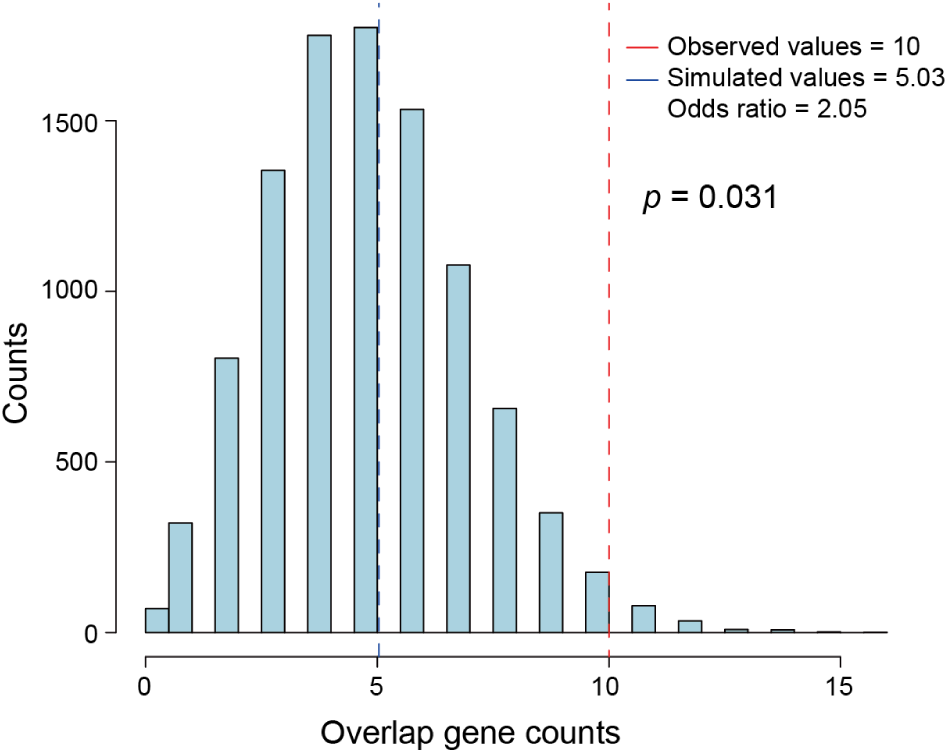
Permutation test for hypoxic relatedness of TESV-associated genes. The null distributions were generated by randomly extracting a set of genes from the entire genome (comprising 27,033 genes) with the same gene counts of hypoxia-related genes (*n* = 473) over 10,000 iterations, and calculated the overlap with the 287 TESV-associated genes. The significance of the difference of the mean value for observed value and expected value were calculated using Chi-square test, and indicated by the *p*-values.

**Supplementary Fig. 11.**
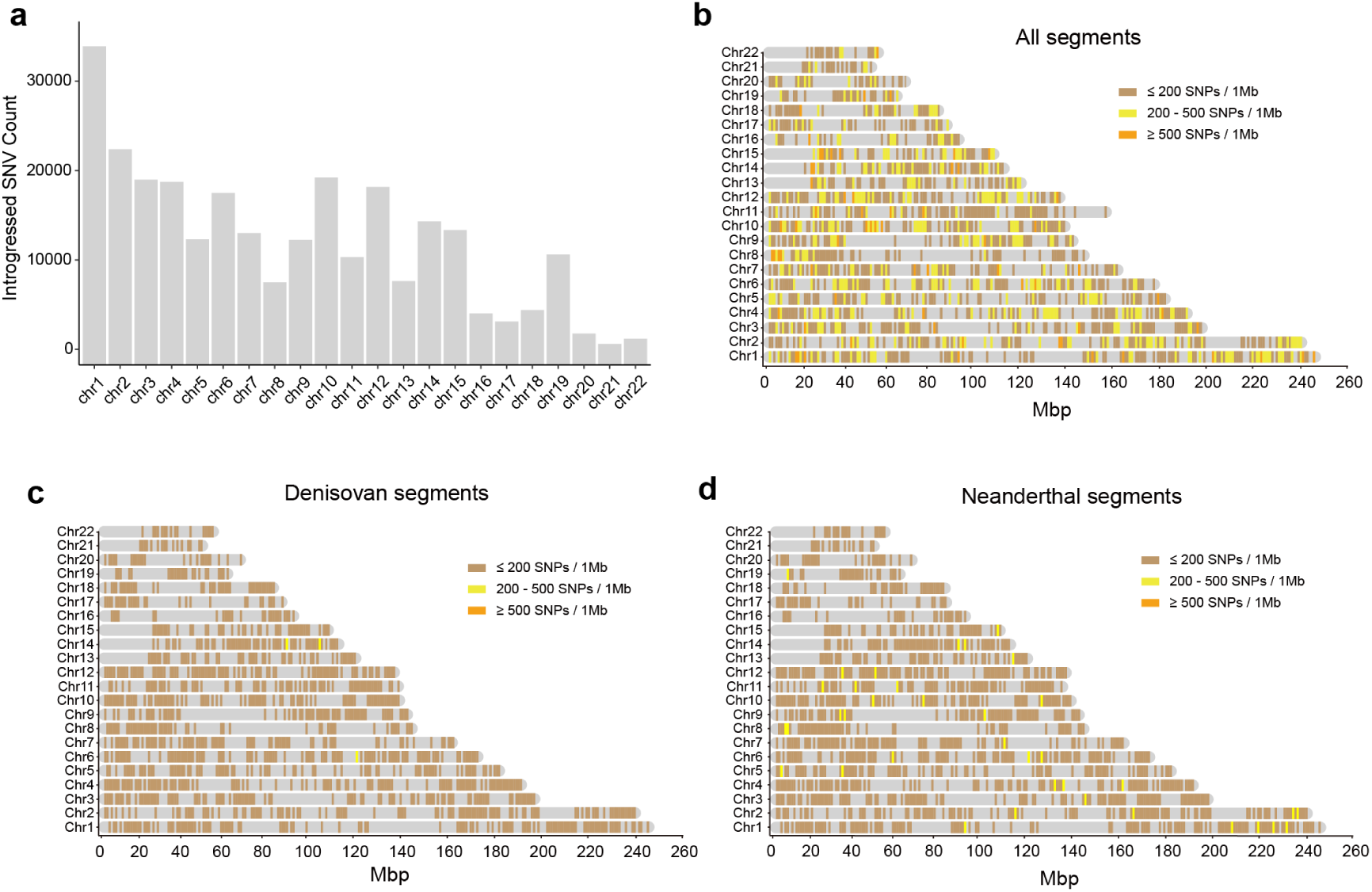
Genome-wide distribution of archaic introgression. **a**, Counts of introgressed SNVs across chromosomes. **b–d**, Heatmaps showing the distribution of all introgressed segments, Denisovan-introgressed segments, and Neanderthal-introgressed segments, respectively, across Tibetan autosomes.

**Supplementary Fig. 12.**
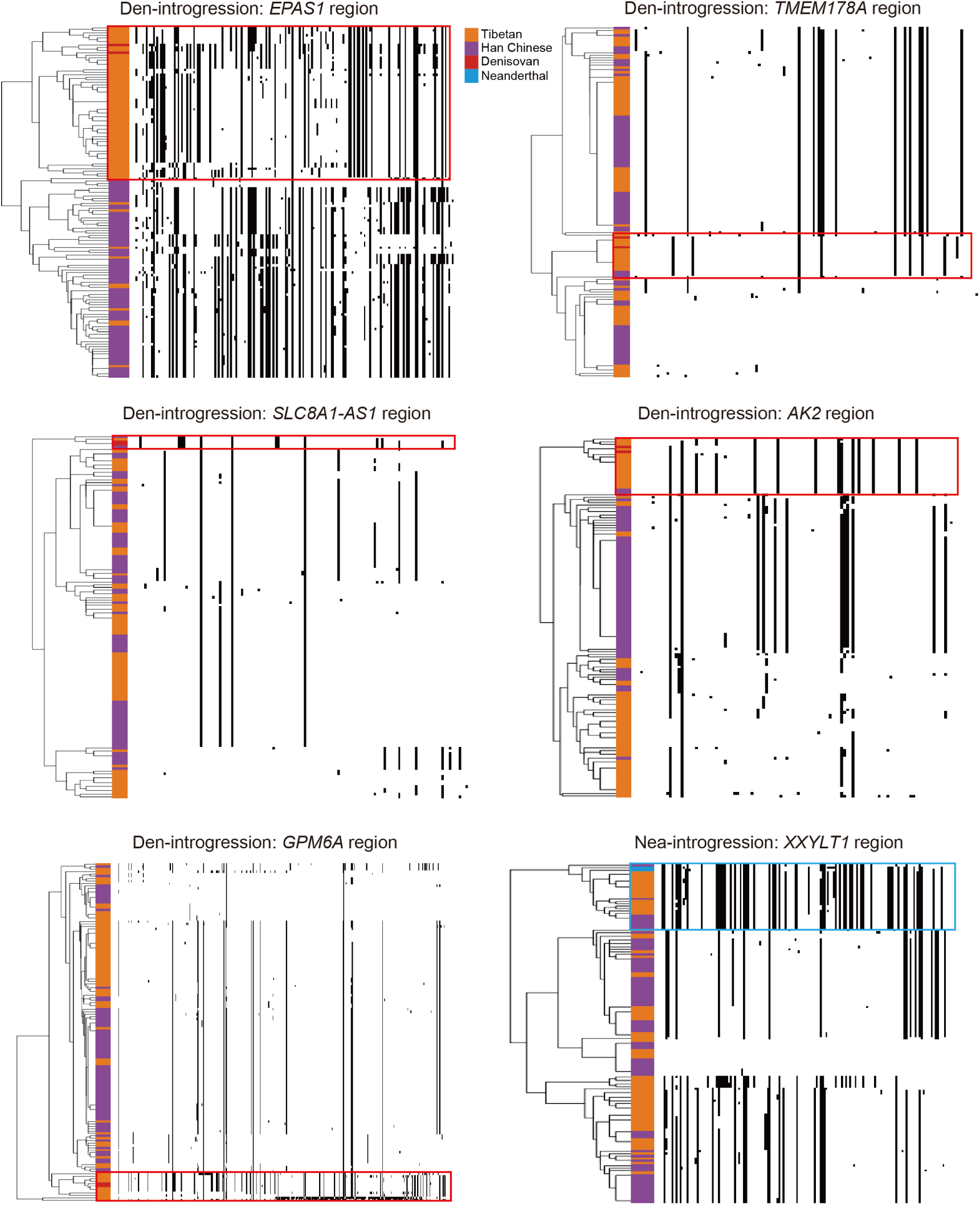
Hierarchical clustering of haplotypes across adaptive introgression regions. Shown are five Denisovan-introgressed signals and one Neanderthal example. Rows represent individual haplotypes from Tibetans (orange), Han Chinese (purple), Denisovan (red), and Neanderthal (blue). Columns mark the genotypes of positively selected variants. Putative archaic haplotypes are outlined in red (Denisovan, Den) and blue (Neanderthal, Nea). Gray and black indicate ancestral and derived alleles, respectively.

